# FrustrAI-Seq: Scaling Local Energetic Frustration to the Protein Sequence Space

**DOI:** 10.64898/2026.02.03.703498

**Authors:** Jan-Philipp Leusch, Miriam Poley-Gil, Miguel Fernandez-Martin, Franco L. Simonetti, Julius Schlensok, Nicola Bordin, Burkhard Rost, R. Gonzalo Parra, Michael Heinzinger

## Abstract

Proteins fold into their native three-dimensional (3D) structures by navigating complex energy landscapes shaped by the biophysical and biochemical properties of their sequence. Once folded, some residues remain locally frustrated, reflecting functional constraints incompatible with optimal packing. This *local energetic frustration* provides important insights into protein function and dynamics, but its analysis typically relies on structure-based energy calculations and remains computationally costly at scale. Here, we introduce an ultra-fast sequence-based prediction of local energetic frustration directly from protein sequences using embeddings from protein language models (pLMs). Our method, coined *FrustrAI-Seq*, enables proteome-wide frustration profiling in minutes (∼ 17 minutes for the entire human proteome on a single NVIDIA H100 GPU) while retaining biologically relevant performance as shown for the *α*-globin and *β*-lactamase family. By eliminating the need for explicit structural or evolutionary information, this approach expands frustration analysis to protein regions and classes that were previously inaccessible, including intrinsically disordered regions and high-throughput *de novo* designed protein datasets. To support reproducibility and large-scale applications, we provide the largest freely available resource of precomputed local frustration scores to date (∼ 10^6^ proteins), along with model weights and complete training and inference code at: github.com/leuschjanphilipp/FrustrAI-Seq.

## Introduction

Proteins are evolved polymers that, unlike random heteropolymers, can reliably and rapidly fold into well-defined three-dimensional structures within biologically relevant timescales, thereby overcoming the Levinthal paradox (1). According to the energy landscape theory and the principle of minimal frustration (2), interactions in the native state are, on average, more favorable than the non-native interactions sampled during folding. This bias yields a funneled energy landscape that efficiently guides the polypeptide chain towards its native state.

Despite this global energetic minimization, not all interactions within the folded protein can be simultaneously optimized. A subset of contacts remains in energetic conflict within the native topology, i.e., they are highly frustrated (Fig. 1a). Typically, up to ∼10% of interactions in globular proteins fall into this category (3) (Fig. 1b), suggesting that residual local frustration is an evolutionarily selected characteristic related to function.

**Fig. 1.**
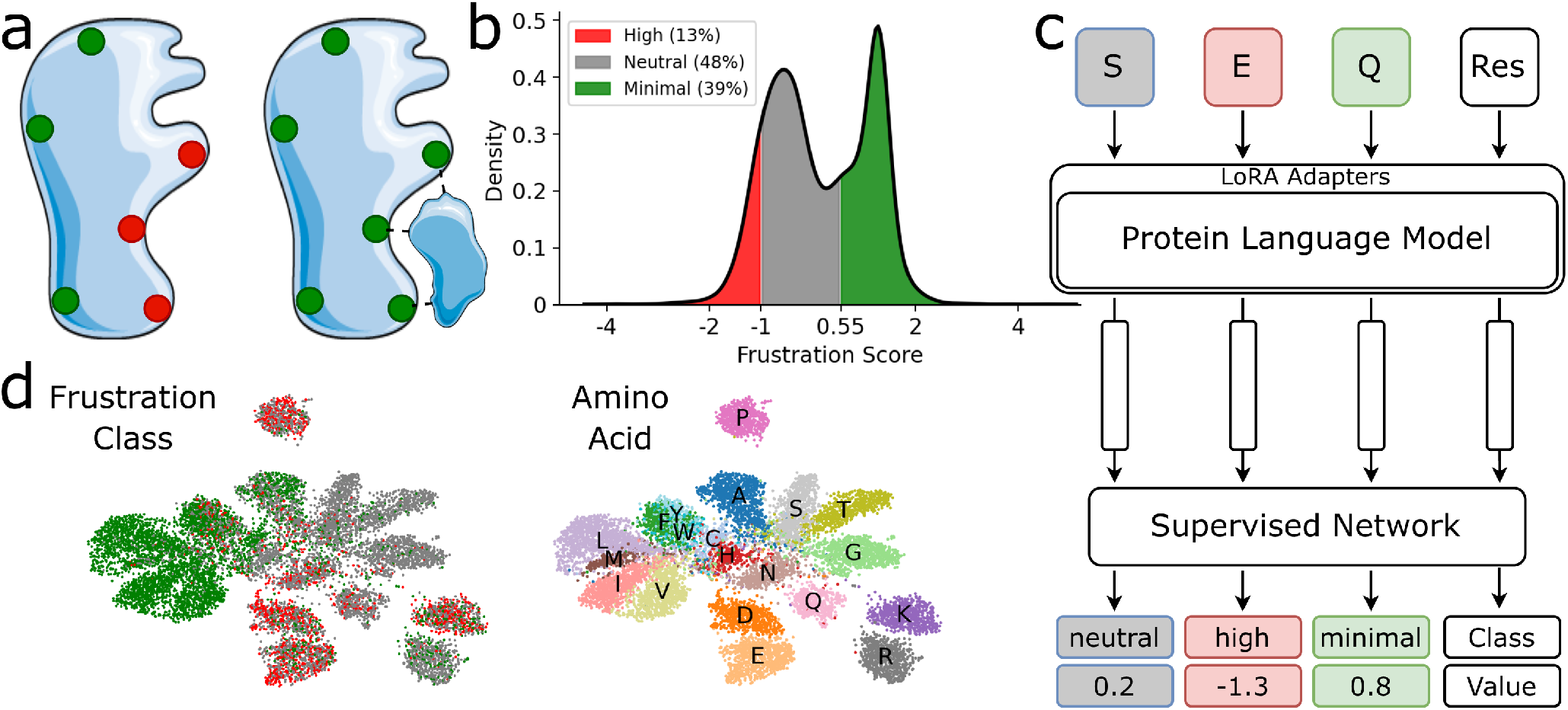
Overview of residue-level energetic frustration and the FrustrAI-Seq model architecture. **(a) Conceptual illustration of local energetic frustration.** Residues in energetically unfavorable, highly frustrated states (left, red) in the unbound form of a protein get stabilized upon ligand binding and become minimally frustrated (right, green). **(b) Distribution of continuous frustration values** obtained from 100 randomly selected training set proteins. Colors indicate the three frustration states: highly frustrated (Frustration Index (FI) < −1.0, 13% of the data), neutral (−1 < FI < 0.55, 48%), and minimally frustrated (FI ≥ 0.55, 39% of the data). **(c) Schematic of the FrustrAI-Seq architecture**. A protein sequence (e.g. “SEQ”) gets embedded by a LoRA fine-tuned protein language model (pLM), here ProtT5. The embeddings are procesed by a shared stack of convolutional layers before being passed to individual prediction heads outputting, for each residue, a continuous frustration score (regression head) and a discrete frustration class (high, neutral, minimal; classification head). **(d) UMAP projection of pLM residue embeddings** from the same 100 random training proteins colored by discrete frustration class (left) and by amino acid identity (right) using ProtT5 before frustration training.

Over the past decade, highly frustrated interactions have been associated with different functional aspects of proteins, such as protein-protein interactions (3, 4), allostery (5), catalysis and co-factors binding (6), and conformational switching (7). These observations suggest that local frustration offers a mechanistic insight into how proteins balance between structural stability and the dynamism required for biological activity (8), with such equilibrium being maintained over evolutionary timescales within protein families (9). Understanding how local frustration shapes the functional aspects of proteins is of ultimate relevance for understanding how changes in their polypeptide sequences can lead to pathological phenotypes (10).

The FrustratometeR, a tool for localizing and quantifying local frustration in protein structures (11, 12) has been used by the community to identify frustration as a key feature underlying several biophysical aspects of proteins (10, 13). The method calculates a frustration index (FI), a z-score obtained by comparing the native energy of a residue or a pairwise interaction with the energy distribution of a set of structural decoys. Based on this FI, residues are classified into one of three frustration states (Fig. 1b): highly frustrated (the native energy is statistically less favourable than the decoys, FI < −1), minimally frustrated (the native energy is statistically more favourable than the decoys, FI ≥ 0.55), or otherwise it is considered neutral (−1 < FI < 0.55). With the advent of novel structure prediction techniques such as AlphaFold2 (14), frustration analysis can be used to study an unprecedented amount of data. However, explicitly calculating frustration, even with a coarse-grained statistical potential as its energy function, becomes computationally costly at large scales, on top of requiring an experimental structure or a predicted model as input, which adds to the computational cost if it needs to be generated.

Large foundational protein language models (pLMs), trained on billions of natural protein sequences, learn representations of the protein space from sequence data alone, either by predicting the next amino acid given all previous ones (auto-regressive language modeling - ALM) (15, 16) or by correcting corrupted input (masked language modeling - MLM) (17–20). While ALM was shown to excel at generating functional proteins (21), MLM models have become the *de facto* standard solution for protein property prediction. First, general-purpose vector representations of protein sequences are generated by extracting the hidden state activations triggered from inputting the sequences to the pre-trained pLM. Those so-called *embeddings* serve as a foundation (sole input) to a second-stage neural network trained specifically to solve a certain task. This strategy of training “experts” on top of general-purpose pLM embeddings has enabled a variety of state-of-the-art predictors, ranging from the prediction of structural features (18) to functional annotations (22), mutational effects (23), binding interfaces (24), and disorder (25, 26). Crucially, pLMs operate at the level of individual proteins and scale efficiently to large datasets.

In this work, we present *FrustrAI-Seq*, a pLM-based method for predicting local energetic frustration at the level of single residues directly from protein sequences. Concretely, FrustrAI-Seq builds on top of the pLM ProtT5 (17) but finetunes the foundation model on a novel dataset of pre-computed frustration values comprising approximately 1 million proteins (Fig. 1c).

Its simplicity and ability to predict frustration without requiring structural models as input enable the method to be applied on the scale of the entire protein universe and be incorporated into high-throughput pipelines aimed at exploring the sequence–frustration landscape, thus improving our understanding of protein evolution and guiding protein design (27). To summarize, our contributions include:

- **FrustrAI-Seq**, an ultra-fast pLM-based method to predict local energetic frustration scores from protein sequences only.
- **The Funstration dataset**, to our knowledge, the largest freely available dataset of local energetic frustration.
- **Confidence** and **discovery scores** for post-hoc filtering of FrustrAI-Seq predictions.
- **Biological relevance and novelty** shown at the level of single proteins, protein families and conformer analysis, zero-shot mutation assessment, frustration– disorder coupling, and large-scale proteome analysis.

## Results

We processed the FunFams (28) dataset from CATH (29, 30) to enrich it with models from TED (31) and curated a subset of high-quality entries to generate the *Funstration dataset*, which constitutes, to our knowledge, the largest freely available pre-computed local energetic frustration resource. This dataset was used to train *FrustrAI-Seq*, a method to predict local energetic frustration at the single-residue level using protein sequence-only input. Towards this end, we fine-tuned the protein language model (pLM) ProtT5 (17) to simultaneously predict continuous frustration scores and discrete frustration states (Fig. 1c).

### 2.1. The Funstration dataset

We introduce the *Funstration dataset* which holds precomputed local energetic frustration values for 186M residues across 983K proteins. We ensure high quality by a) starting from TED domains classified into CATH functional families (FunFams), b) removing highly similar (> 80% sequence identity) sequences, c) keeping only high-quality structural models (median pLDDT score ≥90), and d) avoiding under- or over-representation of very small or large families (restricting family sizes to 20-200). The remaining domains are mapped back to their full protein context, as defined in UniProt, to avoid inputting non-natural protein fragments to the pLM that originate from non-contiguous domain assignments (see Sec. 5.1 for more details).

### 2.2. Local Energetic Frustration Can Be Predicted From Sequence Alone

We used the Funstration dataset (see 2.1), together with low-rank adapters (LoRA) (32), for parameter-efficient fine-tuning of the existing pLM (33), ProtT5, with subsequent shared CNN layers feeding into separate FNN layers to independently predict a) continuous frustration indices (FI) scores and b) binned frustration classes (see 5.1) from protein sequences (see Fig. 1). We evaluated FrustrAI-Seq on a strict hold-out test set, consisting of protein sequences with CATH (34) topologies that the model did not encounter during training (see 5.1). This structure-based split is considerably more stringent than the usual sequence-identity-based splitting yielding comparatively conservative performance estimates. FrustrAI-Seq reaches a Pearson correlation coefficient (*r*) of 0.86 and a mean absolute error (MAE) of 0.33. In comparison, our two baselines (see 5.3) achieve *r* = 0 and MAE= 1.04 (randomly shuffled labels) and *r* = 0.76 and MAE= 0.45 (mean prediction using natural amino acid frustration background). Stratifying per-residue predictions by amino acid type shows that FrustrAI-Seq’s regression head captures overall frustration patterns while under-predicting both long tails (Fig. 2a). This trend is confirmed by comparing the overall FI distributions. FrustrAI-Seq, on average, captures general frustration patterns well, while under-predicting highly and minimally frustrated residues (Fig. 2c). We quantified whether this leads to class confusion by binning the regression output based on FI-thresholds (−1 and 0.55): the minimally frustrated class is predicted reliably (F1 = 0.85, prec. = 0.91, rec. = 0.80), while the highly frustrated class is under-predicted (F1 = 0.54, prec. = 0.76, rec. = 0.42). As reference, our two baselines reach for the highly frustrated class an F1 score of 0.15 (Random) and 0 (Baseline), i.e., a simple mean prediction never predicts high frustration. The recently published FrustraMPNN (35), a fine-tuned ProteinMPNN operating on 3D backbones, reaches a macro F1 of 0.68 on our Funstration test set (see Table 1), confirming that a structure-based encoder captures frustration beyond average amino-acid trends. For the functionally most relevant, highly frustrated class, however, FrustrAI-Seq clearly outperforms it (F1 = 0.54 vs. 0.45), achieving substantially higher precision at identical recall (prec. = 0.76 vs. 0.50 at rec. = 0.42, see Supplementary Table 4). We note that FrustraMPNN was trained on labels computed with the legacy FrustratometeR sequence-separation parameter (SeqDist = 12) rather than the current default of 3, and may have encountered our test proteins during training. We further note that our evaluation was conducted on residues position < 512 as this mirrors FrustrAI-Seqs’ training procedure. An additional test on residues ≥ 512 showed little difference (macro F1 = 0.72, prec. = 0.80, rec. = 0.69, Pearson *r* = 0.85, MAE = 0.35) highlighting FrustrAI-Seq’s ability to generalize not only to unseen porteins during training, but additionally to positions outside the training regime. The class weighting of FrustrAI-Seq’s classification head improves this head’s recall at the expense of precision (F1 = 0.62, prec. = 0.49, rec. = 0.84, see Supplementary Table 5), thereby complementing the regression head. Simply adjusting the regression thresholds has a similar effect, albeit with different dynamics, i.e., at t iso-recall (e.g. rec. = 0.80), the classification head reaches higher precision (prec. = 0.52) than the regression head (prec. = 0.42). To highlight this effect, we computed precision and recall as a function of a) the Shannon entropy of the classification head’s output probabilities and b) continuous regression head output (Fig. 2d). On the held-out test set, correctly classified residues consistently exhibited lower entropy compared to misclassified residues, supporting entropy as a well-calibrated and interpretable proxy for prediction confidence that enables the identification of high-confidence predictions (Fig. 2d).

**Table 1.**
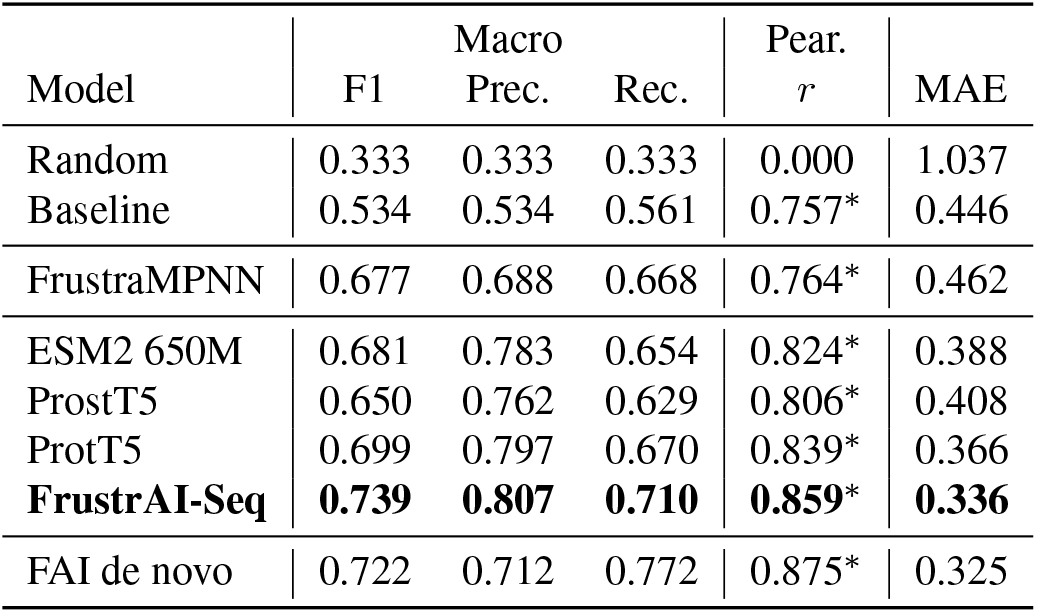
Performance comparison across models. We report key performance metrics for residue level frustration prediction, including macro-averaged F1, precision, recall, Pearson correlation (*r*; * indicates a *p*-value below 0.001), and mean absolute error (MAE) for our strict hold-out test set. The “Random” baseline corresponds to permuted labels, while the “Baseline” uses amino acid type-specific mean frustration and majority class derived from our training data (Fig. 2a) to predict continuous and binned frustration, respectively, for each residue. FrustraMPNN (35) predicts frustration from 3D structures of our hold-out test set using a finetuned ProteinMPNN (36) (see 5.5 for more details.) All other models in the table utilize the same architecture but with different sequence embedding backbones (ESM2-650M, ProstT5, ProtT5). The best-performing pLM, ProtT5, is further fine-tuned with LoRA and class-specific weights, yielding our final model, *FrustrAI-Seq* (FAI). We further report performance for FAI on a hold-out test set of de novo designed proteins (FAI de novo). All numbers reported are derived from the binned regression heads (classification head results, see Supplementary Table 5). Bootstrap standard errors for all metrics are below 0.001 (grouped protein-level bootstrap, 1000 replicates); per-metric values *±* SE are listed in Supplementary Table 4.

**Fig. 2.**
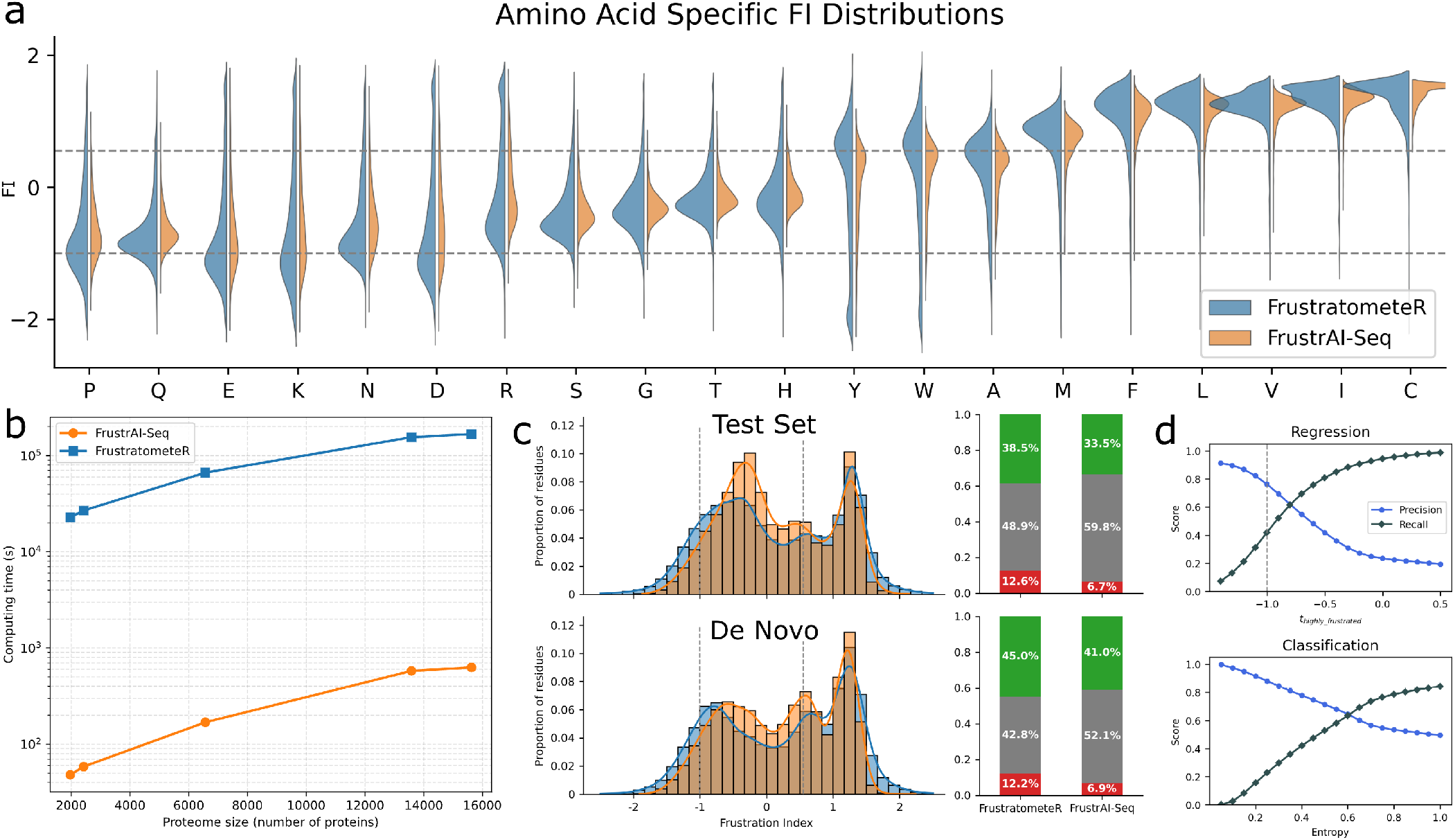
Model output analysis. **(a) Amino acid-specific comparison of FrustratometeR and FrustrAI-Seq predicted frustration.** We compare the FrustratometeR-computed (blue) and FrustrAI-Seq-predicted (orange) frustration indices (FI) individually for the 20 standard amino acids, highlighting amino-acid-specific frustration preferences. Horizontal dotted lines (at -1 and 0.55) indicate the thresholds for the frustration classes. Values are clipped at -2 and 1.55 to minimize the effect of outliers in visualization. **(b) Runtime comparison between FrustratometeR and FrustrAI-Seq**. We show different computation time curves for FrustrAI-Seq and FrustratometeR. FrustrAI-Seq consistently runs roughly three orders of magnitude faster than FrustratometeR across varying proteome sizes. We note that FrustrAI-Seq ran on NVIDIA H100 GPUs, while FrsutratometeR ran on Intel(R) Xeon(R) Platinum 8480 CPUs. **(c) Frustration distribution of FrustratometeR and FrustrAI-Seq**. We compare the FI distributions of FrustratometeR against FrustrAI-Seq’s regression head on our test set (top) and a set of de novo proteins (bottom). Adjacent bar plots show the binned percentages using the default −1 and 0.55 thresholds. **(d) Precision-Recall calibration**. We provide precision (blue) and recall (black) for different regression (top) or classification (bottom) thresholds on predictions of our test set. Classification head thresholds get computed from the entropy over the head’s softmax-normalized output probabilities. At iso-recall of 0.80, the regression and classification heads reach a precision of 0.42 and = 0.52, respectively.

Beyond confidence estimation, we provide for each residue’s predicted frustration a z-score computed with respect to the amino acid-specific mean (our *Baseline* in Table 1) and standard deviation derived from the training set. This normalization enables us to highlight predictions that are outliers within their respective amino acid backgrounds.

### 2.3. Ablation Study

To converge on the final FrustrAI-Seq architecture, we first compared three pLM backbones. ProtT5 performed numerically best on both heads (macro F1 of 0.70 / 0.74 for the regression / classification head, versus 0.68 / 0.73 for ESM2 650M and 0.65 / 0.70 for ProstT5), with the clearest margin on the highly frustrated minority class (regression-head F1 of 0.44, 0.40, and 0.33, respectively). During ablation, performance was assessed on the validation set to not bias our findings on the hold-out test set (see Supplementary Table 1 for full table of ablation results). Finetuning ProtT5 using low-rank adaptors (LoRA) (32) increased performance further for both heads: macro F1 rose by 4 and 3 percentage points for the regression and classification head, Pearson’s *r* increased from 0.84 to 0.87, and MAE decreased from 0.36 to 0.33. To address class imbalance, we applied class weights or focal loss (37) to the cross-entropy loss and evaluated their effects on the highly frustrated class using the classification head. Class weighting reduced precision by 0.19 but increased recall by 0.26, while focal loss reduced precision by 0.19 and increased recall by 0.24, both compared to standard cross-entropy. Neither approach improved the regression head’s binned predictions. Overall, applying class weights resulted in a slight decrease in macro F1 of 0.02 for both cross-entropy and focal loss. Despite a slight decrease in overall performance, we decided to use the weighted cross-entropy for final model training treating this precision–recall tradeoff rather as a feature than a cost. In the highly frustrated class, the regression head is conservative (precision 0.75, recall 0.43 on the validation set), while the class-weighted classification head is permissive (0.49 / 0.85). The confusion matrices in Supplementary Fig. 2 show that these operating points are nested rather than merely different: 99.9% of the residues called highly frustrated by the regression head are also called highly frustrated by the classification head, and no residue is ever assigned to opposite extremes (highly vs. minimally frustrated) by the two heads. The classification head thus returns a superset of highly frustrated calls, and the additional calls are predominantly borderline: 88% of its false positives are neutral rather than minimally frustrated residues (96% for the regression head). We therefore recommend the regression head as the default output, as it reproduces the continuous FrustratometeR index (*r* = 0.86, MAE = 0.33), is threshold-agnostic, and is the more precise of the two when discretized. The classification head is best used as a complementary, recall-oriented filter whenever the objective is to enrich for highly frustrated residues, e.g., when screening large sequence sets for candidate functional sites – and a moderate number of predominantly neutral false positives is acceptable. Taken together, we decided to use the LoRA-tuned ProtT5, trained simultaneously with weighted cross-entropy and regression loss, as the final FrustrAI-Seq model.

### 2.4. FrustrAI-Seq generalizes to de novo designs

The out-of-distribution generalization capability of FrustrAI-Seq was assessed on an additional hold-out test set of 225 de novo-designed proteins derived from the PDA (38) (see 5.7). In general, the de novo set has more minimally frustrated and fewer neutral residues compared to natural sequences (45% vs 38.2% and 42.8% vs 49.7%, respectively; see Fig. 2c) — a trend which is captured well by FrustrAI-Seq (41% vs 32.9% and 52.1% vs 61.2%, respectively). In contrast, the fraction of highly frustrated residues is nearly identical between natural and de novo sequences (12.1% vs 12.2%) — again, a trend which is captured well by our method, albeit with the expected drop in recall (5.9% vs 6.9%). Similar to natural sequences, FrustrAI-Seq tends to under-predict long tails for de novo designs, which mostly affects the recovery of highly frustrated residues while still recovering the vast majority of minimally frustrated residues (rec. = 0.40 vs. rec. = 0.82, respectively). Overall, FrustrAI-Seq captures frustration trends in proteins without evolutionary trace (F1 = 0.86, F1 = 0.78, F1 = 0.52; minimally, neutral and highly frustrated, respectively: see Table 1). When differentiating de novo classes, we observe that designed enzymes follow the natural frustration distribution more closely (delta frustration of 4.1%, 3.2%, 1.0% for minimal, neutral, and high frustration, respectively) than, for example, nanoparticles (8.5%, 10.3%, 1.8%). The same trend is captured by FrustrAI-Seq, at the expected reduced recall of highly frustrated residues (5.8%, 5.8%, 0.0% vs. 10.1%, 12.7%, 2.8%) (see Supplementary Fig. 1 and Supplementary Table 2). To investigate whether FrustrAI-Seq captures frustration in ProteinMPNN designs, we analyzed inverse folded sequences for 21 *α*-globin and 31 *β*-lactamase wild-type structures that were previously generated (39) and found to have less highly frustrated residues than their natural counterparts, and disrupt conserved frustration patterns. Generally, FrustratometeR computed from ESMFold predictions of designed sequences shows high agreement with FrustrAI-Seq predictions for both families (Pearson *r* = 0.91 and *r* = 0.88 for *α*-globins and *β*-lactamases, respectively; Supplementary Fig.4,5).

### 2.5. FrustrAI-Seq captures conserved frustration

Beyond single proteins, we analyzed whether FrustrAI-Seq captures the evolutionary conservation of energetic frustration in protein families. To this end, we compared the Multiple Sequence Frustration Alignment (MSFA) from FrustraEvo (40) to FrustrAI-Seq predictions for a family belonging to one of our hold-out topologies, the *α*-globin family (Fig. 3a).

**Fig. 3.**
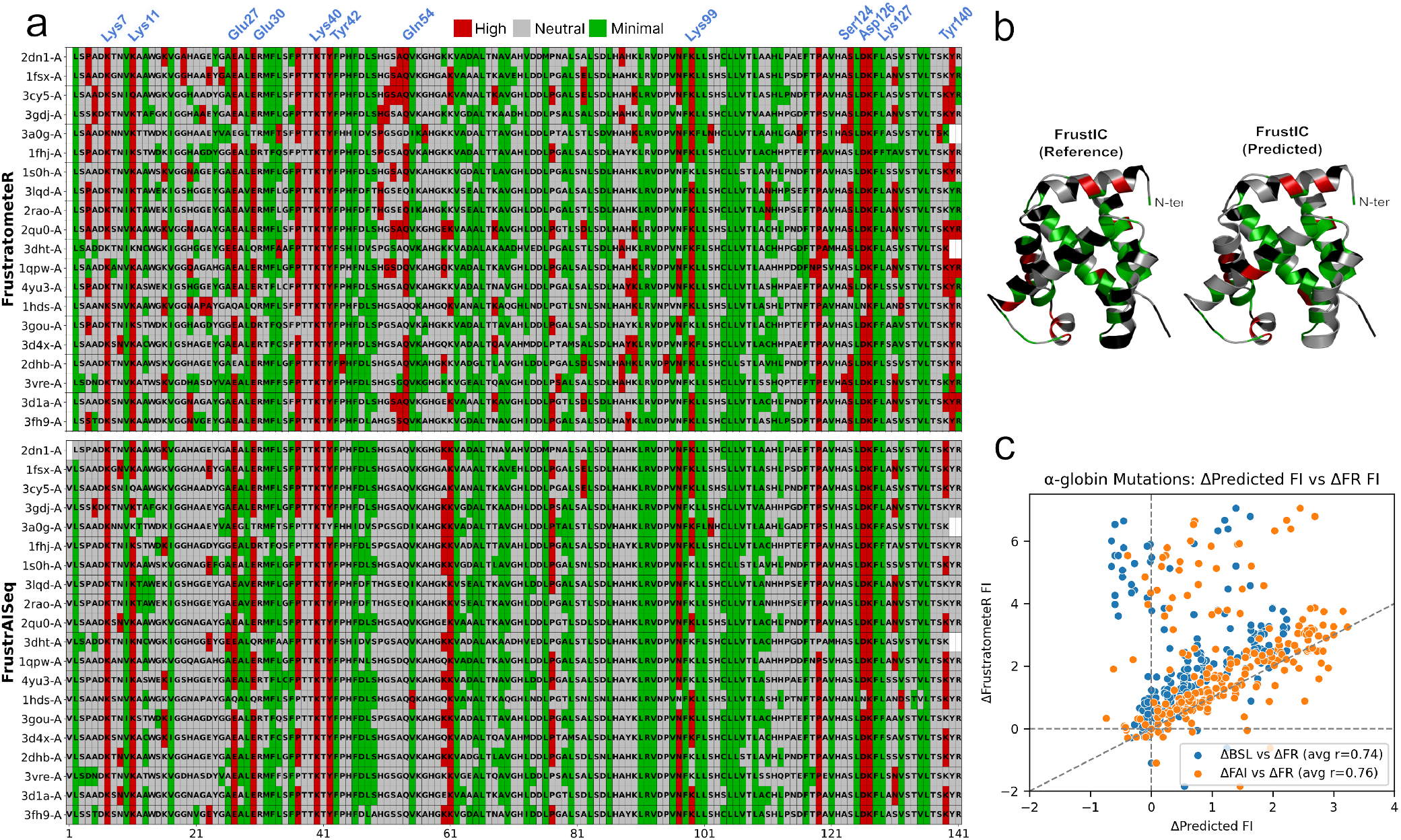
FrustrAI-Seq recapitulates frustration conservation in the *α*-globin family. **(a) Residue-level frustration profiles for the *α*-globin family computed with FrustratometeR and predicted by FrustrAI-Seq.** Heatmaps show residue-wise frustration states across the *α*-globin family for FrustratometeR (top) and FrustrAI-Seq predictions (bottom; frustration states are derived from binned regression head output). Green, red, and gray mark minimally-, and highly frustrated and neutral residues, respectively. Residues with conserved frustration are highlighted in blue at the top. **(b) Structural mapping of Frustration Information Content (FrustIC) on a representative human** *α***-globin structure (2DN1-A)**. FrustIC values derived from FrustratometeR (top) and FrustrAI-Seq (bottom) are projected onto the 3D structure. Residues with FrustIC < 0.5 are shown in black, whereas residues with FrustIC ≥ 0.5 are colored according to the most frequent frustration state in the MSFA. **(c) Recapitualating mutation-induced frustration changes from FrustratometeR with FrustrAI-Seq and *Baseline***. For each of the 12 residues highlighted in (A), we compute the delta frustration between all possible mutants and the native residue (Δ = *FI*_mutant_ − *FI*_native_) with FrustratometeR and compare it against the delta of FrustrAI-Seq (orange, per-site averaged Pearson *r* = 0.76) and our amino-acid-mean Baseline (blue, per-site averaged Pearson *r* = 0.74).

Frustration states predicted by FrustrAI-Seq and computed by FrustratometeR show broad consistency, with conserved patterns of highly frustrated and minimally frustrated residues across homologs associated with functionally important motions (FIMs). Regions where frustration does not agree between methods (columns 4, 50–54, 68, 88–90) are not related to known functional residues of *α*-globins.

We further compared the degree of conservation of frustration states computed from FrustratometeR and predicted by FrustrAI-Seq at each MSFA column by computing a Frustration Information Content score (FrustIC) (9) and projecting them onto a representative human *α*-globin structure (Fig. 3b). FrustIC values produced by both methods show similar conserved frustration patterns, with FrustraEvo-based conservation patterns containing more variable regions. Comparing the FrustIC of both methods shows good agreement (Spearman *ρ* = 0.52).

### 2.6. FrustrAI-Seq recovers conserved frustration from evolutionary ensembles

To better understand the disagreement between FrustratometeR and FrustrAI-Seq within a family, we derived an MSFA for 21 *α*-globins with identical protein sequences but different experimentally measured conformations (conformers). Although functionally relevant columns in the FrustratometeR-derived MSFA share conserved frustration, there are many heterogeneous, non-conserved, highly frustrated, or minimally frustrated residues due to conformational variability (*conformer noise*) in the measured structures (Fig. 4a) originating from biologically unimportant motions (BUMs) (8). Instead, FrustrAI-Seq’s output stays constant as its input does not change. While missing some functionally relevant or conserved positions (e.g. positions 54 or 140), FrustrAI-Seq largely recapitulates conserved frustration patterns while removing conformer noise related to BUMs.

**Fig. 4.**
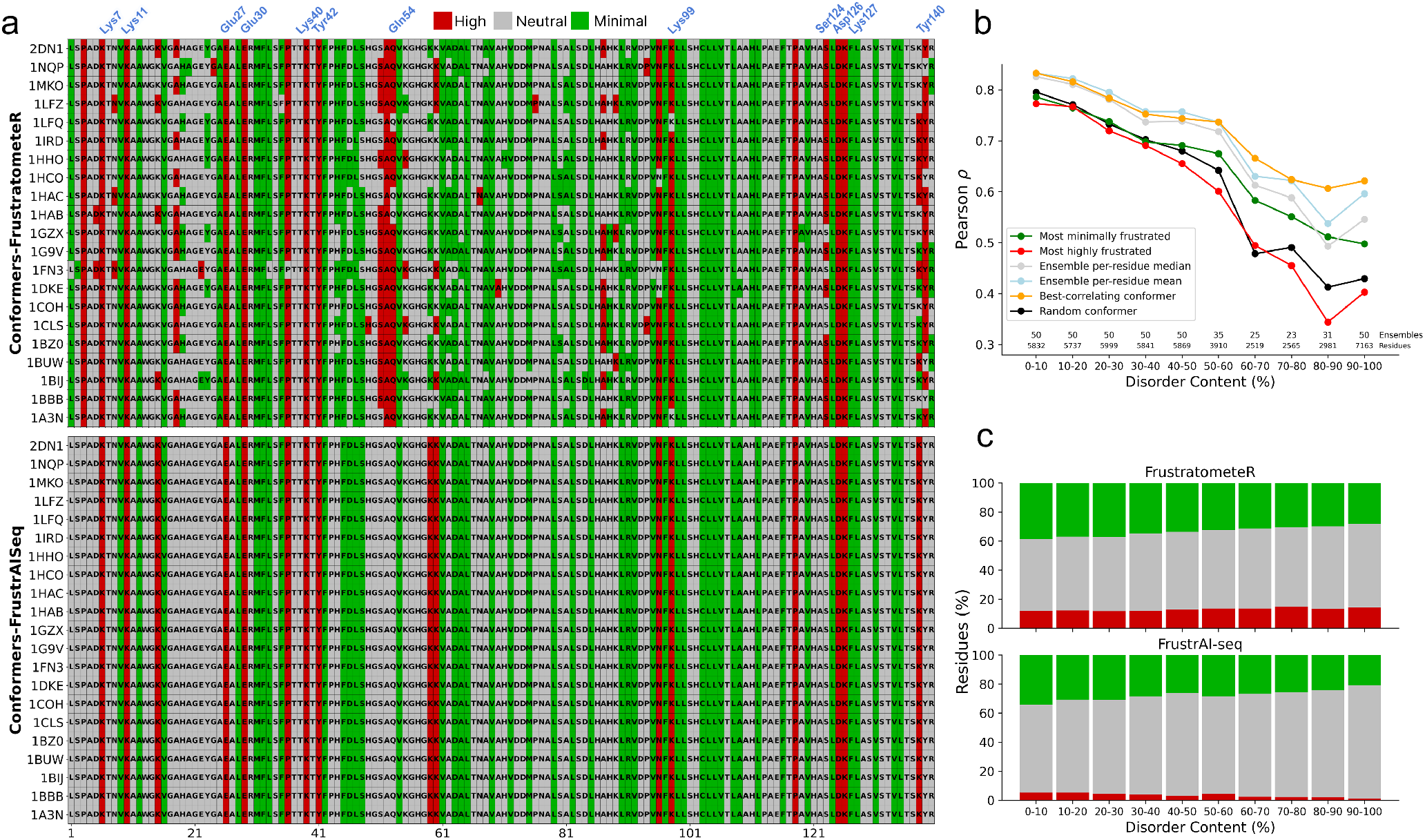
Moving beyond single frustration states: conformer analysis. **(a) FrustrAI-Seq removes conformer noise.** Heatmaps show residue-wise frustration states across different conformers from the Human *α*-globin protein (PDB ID: 2DN1-A) for FrustratometeR (top) and FrustrAI-Seq predictions (bottom; using binned regression output). Red, green, and gray mark highly- and minimally frustrated, and neutral residues, respectively. The 12 natively conserved and highly frustrated residues, mostly involved in PPIs, are labeled in blue at the top. Despite an identical sequence, FrustratometeR produces inconsistent frustration patterns due to different experimental conformations. FrustrAI-Seq instead recapitulates conserved frustration, thereby effectively removing frustration emerging from conformer noise. **(b) FrustrAI-Seq agreement across ensemble representations and disorder content**. Pearson correlation between FrustrAI-Seq predictions and FrustratometeR frustration values over conformational ensembles stratified by disorder content. We report correlations against selected conformers (for most minimally and highly frustrated conformers in each ensemble, and the one conformer with the highest correlation) or residue-level summaries of each conformational ensemble (average and median residue-frustration values over all conformers in an ensemble). FrustrAI-Seq tends to correlate better with the average frustration value of the conformational ensemble, while correlations decrease with higher disorder content. **(c) Frustration state distributions over different disorder content**. Proportion of highly, neutral, and minimally frustrated residues obtained from structure-derived FrustratometeR values (top) and from binned FrustrAI-Seq regression predictions (bottom, thresholds: −1, 0.55), across protein disorder-content bins.

### 2.7. FrustrAI-Seq captures ensemble-averaged frustration across the disorder continuum

Evolutionarily constrained frustration associated with FIMs and the lack thereof associated with BUMs relate to the high-and low-energy states between which a protein oscillates. To analyze on a larger scale whether FrustrAI-Seq is more likely to capture high- or low-energy states, we analyzed 414 proteins, each representing an ensemble of multiple experimentally measured conformers, spanning a total of 9,298 conformers and covering the full order-disorder spectrum. Due to the absence of 3D structure input, FrustrAI-Seq always returns the same frustration for one protein, whereas FrustratometeR frustration is computed for each conformer individually. We report Pearson correlation of both methods for either the conformer with i) lowest or ii) highest overall frustration and put it into perspective of iii) a randomly chosen conformer, the iv) mean and v) median frustration profiles across all conformers in an ensemble as well as vi) the best correlated conformer to FrustrAI-Seq’s output (Fig. 4b). Overall, both methods largely agree for well-structured proteins but increasingly diverge as disorder content increases. Overall, FrustrAI-Seq predictions more closely resemble the ensemble-level mean or median frustration profiles than the frustration pattern of an extreme individual conformer, particularly in highly disordered proteins. Among the extreme conformers, the most highly frustrated consistently shows the lowest correlation, whereas the most minimally frustrated shows the highest. Notably, when considering the best-correlated conformer within each ensemble, the correlation remains above *ρ* ≈0.75 for proteins with up to 50–60% disordered residues and does not fall below *ρ* = 0.6 even for more highly disordered proteins. When comparing the distribution of frustration states along the disorder spectrum, we observed a continuous increase of neutral residues at the expense of minimally frustrated ones for FrustratometeR (Fig. 4c) – a trend that gets captured well by FrustrAI-Seq predictions. As neutral frustration becomes increasingly enriched in more disordered proteins, frustration variability and conformational heterogeneity increase concomitantly (Supplementary Fig.10). We further observe a slight decrease in highly frustrated residues with increasing disorder in FrustrAI-Seq, but not in FrustratometeR. To test whether FrustrAI-Seq embeddings encode robust representations of energetic frustration across the disorder spectrum, we trained a lightweight prediction head on top of embeddings from either FrustrAI-Seq or its base model, ProtT5, to predict disorder as defined by NMR chemical shift scores (41). Following UdonPred (42), we evaluated both models across six disorder definitions, including the latest Critical Assessment of Intrinsic Disorder prediction (CAID3) benchmark (43). FrustrAI-Seq consistently, albeit modestly, outperformed its base model ProtT5 on all benchmarks. The largest improvement was observed when disorder was defined by root-mean-square fluctuation from molecular dynamics simulations (ATLAS test set, Supplementary Fig. 9). To further probe the disorder-frustration relationship, we expanded our comparison to pLDDT, which has been shown to correlate with disorder. However, neither residue-level correlation between pLDDT and frustration, nor local-outlier correlation, i.e. lower local confidence for frustrated residues surrounded by non-frustrated, hinted at any meaningful relationship (Supplementary Fig. 8).

### 2.8. Sensitivity of FrustrAI-Seq to mutations

Within the *α*-globin MSFA (Fig. 3a), we observed that FrustrAI-Seq correctly predicted the frustration of guinea pig *α*-globin (PDB ID: 3A0G chain A) at position 102, which contains a highly frustrated Asparagine (N), despite all other family members harboring a neutral Serine (S) at this position. Following this observation, we systematically tested FrustrAI-Seq’s sensitivity to single amino acid variants (SAVs) at known protein-protein interaction sites in *α*-globins (Fig. 3c, Supplementary Fig. 7) and catalytic residues in *β*-lactamases (Supplementary Fig. 6b-c) to assess the energetic consequences of individual substitutions. Comparison of delta frustration of FrustrAI-Seq between all possible mutants and the native residue (Δ = *FI*_*mutant*_ − *FI*_*native*_) showed a high agreement with FrustratometeR (Pearson’s *r* = 0.76 and 0.88 for *α*-globins and *β*-lactamases, respectively) in both cases. Although FrustrAI-Seq consistently outperformed the *Baseline* model, the amino acid-average predictor achieved Pearson’s *r* = 0.74 and 0.87, respectively.

It has previously been shown that catalytic residues remain conserved and highly frustrated across the majority of *β*-lactamase family members (6) as well as protein-protein interaction sites in *α*-globins (9). FrustrAI-Seq recapitulates these conserved frustration patterns directly from sequence (Supplementary Fig. 3 and Supplementary Fig. 6a), indicating that evolutionarily preserved energetic constraints can be recovered without structural information.

FrustrAI-Seq and FrustratometeR showed strong agreement across the mutational space of catalytic residues, with Pearson correlations ranging from 0.70 to 0.97 across native and mutant variants (see Supplementary Fig. 6b). The highest correlation was observed for the highly conserved and highly frustrated lysine (*rho* = 0.97). Mutation-induced changes in local energetic frustration were also consistent between the two methods (see Supplementary Fig. 6c), with mutations producing net positive frustration changes showing higher correlations than those producing net negative changes, which occurred predominantly at residues S70, S130, and A237. Finally, both methods predicted that numerous single amino acid decrease frustration compared to the native catalytic identities (Supplementary Fig. 6d), consistent with previous analyses in enzymatic families (6).

Similarly, FrustrAI-Seq and FrustratometeR showed strong agreement across the mutational landscape of *α*-globin protein-protein interaction sites, with Pearson correlations ranging from -0.03 to 0.98 (mean Pearson *r* = 0.76) (see Fig. 3c and Supplementary Fig. 7). The highest agreement was observed for residues K7, E30, K40 S124, D126 and K127, whereas lower correlations were restricted to a small number of positions (K11, Q54, and Y140). Overall, these results indicate that FrustrAI-Seq accurately captures mutation-induced changes in local energetic frustration at conserved protein-protein interaction sites directly from sequence.

### 2.9. Local frustration analysis at unprecedented high--throughput scale

To illustrate the speed and scalability of FrustrAI-Seq, we inferred residue-level frustration profiles for ≈ 136 million proteins contained in 22,105 UniProt reference proteomes.

Where previous structure-based analysis showed that well-folded monomeric proteins contain ∼10% highly frustrated-, 40% minimally frustrated-, and 50% neutral residues (3, 10), our proteome-scale analysis found, among the 5,000 proteins with the highest proportion of each frustration class, 20.9– 29.1% highly-, 61.5–98.7% minimally frustrated, and 88.8–100% neutral residues.

The highest proportion of highly frustrated residues was found in a 97-residue uncharacterized protein from Solanum tuberosum (UniProt ID: M1DY82), with 30.9% of its residues classified as highly frustrated and showing high agreement between FrustrAI-seq and FrustratometeR Supplementary Fig. 11a). Proteins with the highest proportions of minimally frustrated residues were predominantly homopolymers or contained extensive low-pLDDT regions (Supplementary Fig. 11b). In contrast, proteins with the highest proportions of neutral frustration showed poor agreement between FrustrAI-Seq and FrustratometeR, and most correspond to entries with predicted or uncertain evidence in UniProt. Summary statistics (frustration state proportions, GO enrichments for each subset, and representative FrustrAI-Seq/FrustratometeR comparisons) from this 136-million protein dataset can be found in Supplementary Fig. 11).

When restricting the analysis to high-confidence proteins (protein- or transcript-level evidence; n = 255,794), the distribution of neutral frustration remained largely unchanged, whereas the highly and minimally frustrated classes became less extreme. In what follows we describe what we find in the top 2,500 proteins from each class. The maximum proportion of highly frustrated residues decreased to 20.9%, although the top-ranking proteins are predominantly transmembrane proteins. Transmembrane regions are usually enriched in high frustration because the AWSEM-MD Hamiltonian (44) used by FrustratometeR assumes an aqueous environment rather than a lipid bilayer. As a result, both FrustratometeR and FrustrAI-seq can identify putative transmembrane regions, whereas a membrane-aware Hamiltonian would instead be expected to classify these residues as minimally frustrated (45).

Excluding transmembrane proteins, the three most highly frustrated proteins were an FAD assembly factor SdhE from *Vibrio cholerae serotype O1* (86 residues; 18.6% highly frustrated, UniprotID Q9KPA2) (Fig. 5b top), a MarR-family transcriptional regulator from *Saccharolobus solfataricus* (144 residues; 17.3%, Uniprot ID Q97YG9), and the MIP18 family-like domain-containing protein from *Bacillus anthracis* (105 residues, 16.2%, Uniprot ID A0A6L8PPP9). At the opposite extreme, the most minimally frustrated protein is NADH-ubiquinone oxidoreductase chain 4 from *Bos taurus* (409 residues, 60%, Uniprot ID P03910) (Fig. 5b middle) . Lastly, numerous proteins contained nearly 100% neutral residues. Inspection of their available UniProt structural models revealed that most correspond to low-confidence predictions (extensive unstructured regions and low pLDDT scores), suggesting they are intrinsically disordered proteins. A smaller subset displayed plausible structures despite uniformly low pLDDT values. In these cases, FrustrAI-Seq predicted predominantly neutral frustration, whereas FrustratometeR produced frustration profiles resembling those of well-folded proteins. One example is the putative uncharacterized protein LOC400692 from Homo sapiens (123 residues; 80% neutral, Uniprot ID Q8N3U1) (Fig. 5b bottom). Although the AlphaFold2 model adopts a compact fold, the uniformly low pLDDT scores suggest low structural confidence. Consistently, MobiDB (46) classifies this protein as highly disordered, raising the possibility that the predicted fold represents a conditionally stabilized state, for example upon binding to a partner or under specific cellular conditions.

**Fig. 5.**
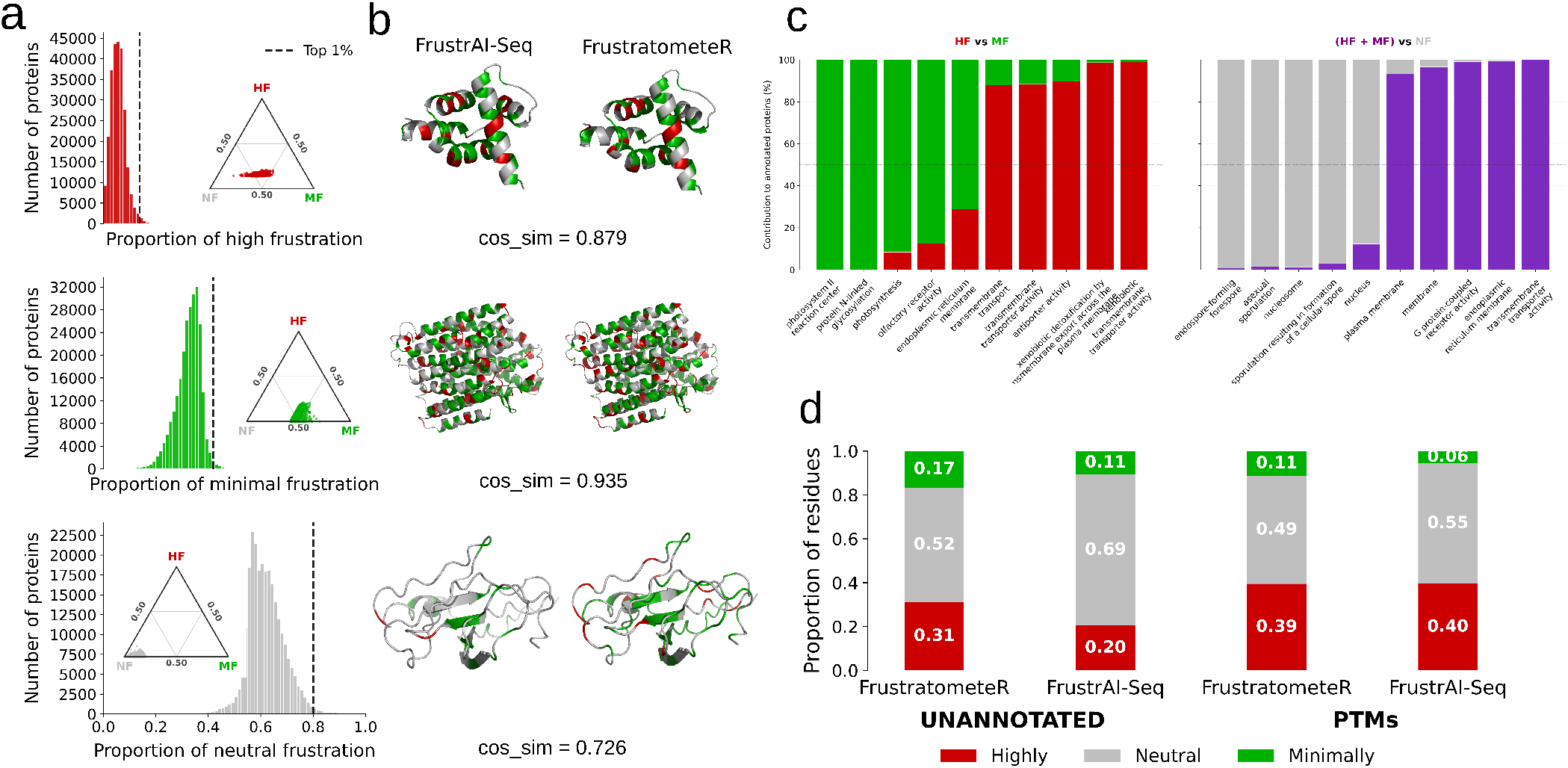
Proteome-scale Frustration Analyses. **(a)** Distributions of the per-protein proportions of highly frustrated (top), minimally frustrated (middle), and neutral residues (bottom) across 255,000 proteins from 22,000 proteomes supported by both protein- and transcript-level evidence. Dashed vertical lines indicate the top 1% of proteins for each frustration class. Insets show ternary representations of the same top 1% proteins according to their relative proportions of highly frustrated (HF), minimally frustrated (MF), and neutral (NF) residues. **b)** Representative proteins selected from the top 1% of each frustration class, comparing residue-level frustration assignments obtained with FrustrAI-Seq (left) and FrustratometeR (right). Residues are colored according to frustration class using frustration values calculated by FrustratometeR or predicted by FrustrAI-Seq. Cosine similarity values quantify the agreement between the two residue-level assignments. From top to bottom: Q9KPA2, P03910 and Q8N3U1. **c)** Left, comparison of GO terms enriched among the top highly frustrated (HF) and top minimally frustrated (MF) proteins. Right, comparison of GO terms enriched among proteins with high proportions of either highly or minimally frustrated residues (HF + MF) and proteins with high proportions of neutral residues (NF). Bars indicate the relative contribution of each protein group to the enriched GO terms. **d)** Comparison of frustration-class proportions between post-translationally modified (PTM) residues and an amino-acid-composition-matched set of unannotated residues. Residues were classified as highly frustrated, neutral, or minimally frustrated (green) according to either structure-based FrustratometeR calculations or FrustrAI-Seq regression predictions.

Functional characterization of this high-confidence subset via gene ontology (GO) enrichment analysis revealed clear differences between proteins with extreme frustration compositions (Fig. 5c): while highly frustrated proteins were strongly associated with membrane-related processes, including trans-membrane transport and transporter activity, minimally frustrated proteins were preferentially associated with intracellular and metabolic functions. By contrast, mostly neutral proteins were generally shorter and enriched in nuclear and gene regulatory functions, including DNA binding, transcription factor activity, chromatin organization, and regulation of transcription by RNA polymerase II. Representative comparisons of those extreme cases with FrustratometeR showed good agreement for highly and minimally frustrated proteins, whereas mostly neutral proteins displayed substantially lower agreement.

To investigate whether post-translationally modified residues exhibit distinct energetic properties, we compared the frustration states of 7,139 UniProt-annotated modified residues from the Funstration dataset with an equal number of unannotated residues (following the distribution of amino acid identities). Both FrustratometeR and FrustrAI-Seq showed a marked enrichment of highly frustrated residues at modification sites, accompanied by a depletion of minimally frustrated residues (Fig. 5d). While FrustratometeR classified 39% of modified residues as highly frustrated compared to 31% of unannotated residues, FrustrAI-Seq increased this contrast from 20% to 40%, indicating a stronger separation between modified and unmodified positions.

These analysis are only possible because FrustrAI-Seq scales efficiently from individual proteins to whole proteomes (Fig. 2b). As a reference, computing the frustration values for the entire human proteome, using the structural models from AlphaFoldDB (47), takes about 17 minutes on a single Nvidia H100 GPU and 58 hours with FrustratometeR without considering the prediction of structures.

## Discussion

Energetic frustration is a central concept in protein bio-physics, linking folding, dynamics, and function (48). However, frustration analysis has traditionally required experimental structures or high-quality structural models and computationally intensive energy calculations, limiting its applicability to large-scale and exploratory studies.

In this work, we release the *Funstration dataset*, a collection of high-quality functional protein families, constituting the largest set of pre-computed residue-level frustration scores to date, comprising approximately one million proteins (Sec. 2.1). Using this dataset, we trained *FrustrAI-Seq*, a protein language model (pLM)-based frustration prediction method, showing that residue-level energetic frustration can be inferred directly from protein sequence without explicit structural input. Compared with previous methods, FrustrAI-Seq provides a three-order-of-magnitude speed-up (Fig. 2b), enabling proteome-scale frustration annotation (17 min for the human proteome on a single NVIDIA H100 GPU; Sec. 2.9), while preserving biologically meaningful frustration patterns across individual proteins (Sec. 2.2), protein families (Sec. 2.5), mutational perturbations (Sec. 2.8), and *de novo* designed proteins (Sec. 2.4).

The ability of FrustrAI-Seq to generalize to *de novo* designed proteins, which lie outside natural evolutionary sequence space (and were not used for training), addresses a fundamental question: can sequence alone encode sufficient information to recover local energetic frustration, a property formally defined for a specific three-dimensional protein conformation?

To better support our affirmative answer to this question, we first stress that all reported performance was measured on a strict hold-out test set whose CATH topologies were entirely excluded from training; second, FrustrAI-Seq clearly outperformed a *Baseline* approach predicting average amino acid-type frustration means; third, crucially, this same *Baseline* never predicts any residue to be highly frustrated, as no mean was below the high frustration threshold; fourth, our method assigns different frustration states to near-identical local sequence contexts. Together, these results indicate that FrustrAI-Seq not only recovers frustration signatures beyond simple average trends but also generalizes to folds not seen during training.

Importantly, while pLMs operate on sequence-only input, these models have been shown to implicitly learn aspects of structure during their self-supervised pre-training (cite). Recent interpretability studies have also shown that pLMs primarily store coevolutionary coupling statistics, analogous to Potts models rather than explicitly learning folding physics, consistent with evolutionary constraints being embedded in sequence embeddings (49). The energy landscape theory predicts that evolutionary and physical energy functions are correlated, implying that sequence statistics implicitly encode energetic constraints on folding and function (50). FrustrAI-Seq leverages this correlation by explicitly aligning sequence embeddings with physics-derived energetic frustration, effectively projecting features of the physical energy landscape into evolutionary latent space. Frustration therefore remains a structure-dependent property, but the structural and evolutionary information required to approximate it is recoverable from sequence through pLMs.

However, the dynamic nature of proteins gives rise to multiple frustration signatures per protein, one per conformer. This raises the question: in the absence of explicit structure input, which frustration profile is approximated by FrustrAI-Seq?

By analyzing frustration patterns across conformational ensembles spanning different degrees of sequence disorder, we found that FrustrAI-Seq (a) preferentially captures the energetic frustration profile from the conformer that is closest to the ensemble-average, (b) agrees closely with FrustratometeR for highly ordered proteins, whose conformers share similar structures and energies, and (c) gradually loses correlation as disorder increases, although the best-correlated conformers still reach *ρ* > 0.75 for proteins with up to 50–60% disorder and remain above *ρ* = 0.6 even for more highly disordered cases. These findings are especially noteworthy given that we excluded AlphaFold2 predictions with an average pLDDT below 90 during data pre-processing, effectively removing proteins with substantial intrinsically disordered regions (51). Furthermore, this trend mirrors the results obtained from conformational ensembles using FrustratometeR (Fig. 4c) and agrees with the weakly funneled energy landscapes proposed for intrinsically disordered proteins (52). The ability of FrustrAI-Seq embeddings to improve disorder prediction relative to ProtT5 embeddings further supports a close relationship between local energetic frustration and intrinsic disorder, suggesting that both properties emerge from common sequence-derived biophysical constraints (Supplementary Fig. 9).

Having established that FrustrAI-Seq is able to reliably recover energetic frustration directly from sequence, we next consider how this capability enables evolutionary, mutational, and proteome-scale studies that were previously impractical or impossible.

By not relying on protein structures as input, FrustrAI-Seq is independent of both structural model quality and biologically unimportant motions (BUMs) (8) resulting in conformer noise which together obfuscate the evolutionarily constrained frustration associated with functionally important motions (FIMs). Distinguishing these two components was the original motivation behind FrustraEvo (9, 40), which identifies frustration patterns conserved across homologous proteins to separate evolutionarily constrained energetic signals from conformer-specific fluctuations.

Consistent with previous structure-based FrustraEvo analysis in the *α*-globin family (9), FrustrAI-Seq recovers the conserved frustration signatures associated with protein function and stability. Likewise, when analyzing multiple experimental conformations of human *α*-globin, where the sequence remains fixed and frustration differences arise solely from conformational variability, FrustrAI-Seq preserves conserved functional frustration patterns while smoothing conformer-specific fluctuations. Beyond providing biological insight, this substantially extends the high-throughput applicability of FrustraEvo to protein family resources such as CATH (53), Pfam (54), and other large sequence databases.

The same principle also extends to mutational analyses. During the *α*-globin analyses, we observed that individual amino acid substitutions produced distinct frustration predictions despite nearly identical local sequence contexts (e.g. position 102 in Fig. 3a), suggesting that FrustrAI-Seq captures sequence constraints extending beyond the immediate amino acid environment and consistent with long-range evolutionary couplings. This observation was supported by the strong agreement between mutation-induced frustration changes predicted by FrustrAI-Seq and FrustratometeR for both protein-protein interfaces in *α*-globins and catalytic residues in *β*-lactamases. Although the Baseline model (amino-acid-specific average frustration values) explained much of the variation at these highly conserved functional sites, FrustrAI-Seq consistently performed better, indicating that it captures context-dependent energetic effects beyond residue identity alone. The strong Baseline performance may be inflated as we biased our analysis towards catalytic residues which show very distinctive frustration profiles across homologs, with many of them conserved and highly frustrated (Fig. 4a), suggesting that energetic constraints are evolutionarily preserved at functionally critical sites. Moreover, whereas FrustratometeR estimates mutational frustration for a single structural conformation, FrustrAI-Seq predicts frustration directly from sequence and may therefore capture the conformationally robust component of mutational effects. Future work combining ensemble-based structural methods such as BioEmu (55) with largescale experimental resources such as ProteinGym (56) will help determine to what extent sequence-based frustration prediction can improve variant effect prediction and protein engineering applications.

Beyond natural proteins, FrustrAI-Seq also generalizes to *de novo* designed sequences, despite these lacking the evolutionary history pLMs are trained on. Across multiple classes of *de novo* proteins, FrustrAI-Seq accurately recapitulates the characteristic frustration distributions obtained from structure-based calculations, indicating that its predictions extend well beyond natural sequence space (Supplementary Fig. 1). Similarly, the strong agreement between FrustrAI-Seq and FrustratometeR for inverse-folded Protein-MPNN designs demonstrates that the learned frustration representations remain applicable to computationally designed proteins (see Supplementary Figure 4a and 5a). Interestingly, FrustrAI-Seq also captures the systematic reduction in local energetic frustration introduced by inverse folding as designs are not conditioned to satisfy functional constraints. Preserving native highly frustrated regions in reverse folding designs could therefore represent an additional filtering step in design pipelines to improve functional hit rates in protein engineering and avoid loss of function as in (57).

Due to its speed and scalability, FrustrAI-Seq extends local frustration analysis from individual proteins to complete proteomes. This enabled the first survey of local energetic frustration across approximately 136 million proteins, from 22,105 proteomes present in Uniprot. We have seen that the upper limit for high frustration is for transmembrane proteins having ≈30% and for soluble proteins ≈18%, for minimally frustrated regions ≈60% and for neutral it increases to maximums in disordered proteins. In terms of Gene Ontology terms enrichment, proteins enriched in highly frustrated residues were predominantly associated with membrane-related functions, whereas minimally frustrated proteins were enriched in intracellular and metabolic processes. In contrast, proteins with high proportions of neutral frustration were enriched in nuclear and gene regulatory functions and frequently corresponded to intrinsically disordered proteins, suggesting that sequence-based frustration predictions may better reflect their underlying conformational ensembles than frustration estimates derived from a single structural model. Deeper exploration of this unprecedented dataset will facilitate the study of how local energetic frustration emerges from protein sequences across the folded and disordered regions of the protein universe and its relationship to protein function.

The structure-agnostic nature of FrustrAI-Seq also defines limitations to its applicability. Because single predictions are derived from each sequence, FrustrAI-Seq cannot resolve frustration changes associated with structural transitions, molecular dynamics trajectories, ligand binding, or allosteric conformational changes. Structure-aware methods such as FrustratometeR and FrustraMPNN therefore remain the appropriate tools for studying systems in which distinct conformational states are characterized by different frustration patterns, such as the open and closed conformations of adenylate kinase (58). Similarly, variant effect analysis may require evaluating the impact of mutations on specific conformational states of a protein. Once structure prediction methods are capable of capturing such effects, structure-based frustration methods remain better suited.

Another potential limitation inherited from the underlying pLM is that prediction quality may depend on how well a given sequence is represented in the learned embedding space. Previous studies have shown that the performance of pLMs in downstream tasks such as protein structure prediction (59, 60) and variant effect prediction (56) is influenced, at least in part, by the amount of evolutionary information available for a sequence. Whether similar biases affect sequence-based frustration prediction remains to be systematically evaluated, particularly for proteins occupying sparsely sampled regions of sequence space. Good agreement between FrustratometeR and FrustrAI-Seq on de novo-designed sequence and ProteinMPNN designs, which, by definition, lack any evolutionary history, make us confident that our method still performs reasonably well in such cases.

More broadly, robust training of conformation-aware frustration models will require paired datasets capturing conformational variability together with frustration annotations, allowing models to account for multiple valid conformational states. Future work could also incorporate side-chain information, post-translational modifications, non-standard amino acids, or lipid layers for frustration calculation, but currently FrustratometeR’s coarse-grained force field is also blind to such changes.

Importantly, FrustrAI-Seq complements physics-based FrustratometeR calculations, which remain the reference framework for energetic frustration analysis, providing more accurate structure-based estimates with configurational and mutational frustration calculations at the level of contact maps. Together, FrustratometeR and deep learning-powered tools such as FrustrAI-Seq and FrustraMPNN establish a multiscale framework that integrates physical energy landscape theory with evolutionary sequence representations, enabling frustration analysis from individual structures to proteome-scale and synthetic sequence space produced by generative AI models.

FrustrAI-Seq establishes local energetic frustration as an AI-inferable evolutionary constraint and provides a scalable framework for integrating energy landscape theory into sequence-based models of protein biophysics, enabling frustration analyses at the scale of the protein universe while also serving as a transferable representation for next-generation AI pipelines, as recently demonstrated by its application to improve conformational sampling from multiple sequence alignments (61).

## Supporting information

FrustrAI-Seq_Supplementary

## Availability

FrustrAI-Seq is available at github.com/leuschjanphilipp/FrustrAI-Seq. The associated training data (Funstration dataset) and model weights are available at huggingface.co/datasets/leuschj/Funstration and huggingface.co/leuschj/FrustrAI-Seq.

## Methods

### 5.1. The Funstration dataset

Our dataset is derived from a large-scale collection of protein domains classified into CATH functional families (62), which provide functionally coherent groupings of homologous domains within CATH superfamilies (34). FunFams were defined using the FunFH-MMer protocol (63), which partitions domain superfamilies based on conserved and specificity-determining positions. Protein domain sequences were clustered at 80% sequence identity using CD-HIT (64, 65), retaining one representative per cluster and restricting family sizes to 20-200 sequences, thereby ensuring the capture of relevant evolutionary and energetic signals. Such filtering also prevents over-representation (and thus overfitting) of a few but very large CATH superfamilies. The corresponding protein 3D structures were obtained from the PDB (66, 67) and TED (31) databases. Prior to frustration analysis, domains were filtered using structural quality criteria: high-confidence predicted structures (predicted Local Distance Difference Test (pLDDT) ≥ 90), sufficient secondary structure content (Secondary Structure Elements (SSE) > 6), and globularity metrics (packing density > 10.333 and normed radius of gyration < 0.356). After filtering, the final dataset comprised 983,212 protein domain sequences, totaling 186,477,919 residues and spanning 8,259 FunFams.

Using the FrustratometeR algorithm (3, 68), single-residue local frustration was quantified for all residues by comparing the energetic contribution of the native amino acid at each position to a distribution of energies obtained from alternative amino acid substitutions. The resulting continuous frustration index ranges from −6.94 to 8.67. The distribution was centered near zero (median = 0.05; IQR: −0.66 to 1.09), with 95% of values falling between −1.50 and 1.66. Following (3, 9), frustration indices were discretized to classify positions as highly frustrated (frustration index ≤−1), neutral (−1 < frustration index < 0.55), or minimally frustrated (frustration index ≥0.55).

To ensure that protein language models receive full contextual information, we mapped each domain subsequence to its corresponding full-length protein sequence using UniProt (2025-03) (69) and AlphafoldDB (v4) (70). This step mitigates the risk of presenting out-of-distribution samples to our pLMs, i.e., non-natural proteins resulting from non-contiguous domains.

To rigorously evaluate model generalization, we split the dataset into train, validation, and test sets based on CATH topology assignment, ensuring that no topology appears in more than one set. This splitting strategy prevents data leakage of similar structures across sets, which could otherwise inflate performance metrics and hinder generalization. Additionally, we ensured that none of the CATH topologies associated with the use-case proteins were present in the training set. After splitting, the training set contained 887k sequences from 499 distinct CATH topologies and 1,083 homologous superfamilies. The validation set contained 49k sequences from 30 topologies and 61 superfamilies. The test set contained 48k sequences from 33 topologies and 56 superfamilies. In agreement with previous studies (3), 12.5% of the residues are highly frustrated, 48.4% are neutral, and 39.1% are minimally frustrated. These proportions are consistent across the train, validation, and test sets. We dub this filtered, pre-split dataset, the *Funstration dataset*.

### 5.2. FrustrAI-Seq architecture and training

The proposed FrustrAI-Seq takes as input a protein sequence *X* and predicts for each residue *r*_*i*_ of that protein sequence its local energetic frustration *y*_*i*_. Let *X* = *r*_1_, *r*_2_, …, *r*_*L*_ be a protein sequence of length *L*, where each residue *r*_*i*_ ∈ *AA* = *A, C*,…, *Y* is from the set of the 20 canonical amino acids (AA). 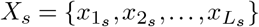 denote the FrustratometeR-derived frustration values, whereas 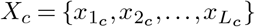 denote the ground-truth class labels. FrustrAI-Seq then simultaneously outputs a continuous frustration score *y*_*i*_*s* and a discrete frustration class 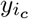.

FrustrAI-Seq comprises two major components: a protein language model (pLM) and a supervised head. For the pLM, we chose the encoder of ProtT5 (specifically, the encoder part of ProtT5-XL-U50, as described in the original publication (17)), a pre-trained pLM trained using masked language modeling on billions of protein sequences. To balance computational efficiency with sequence coverage, we applied a maximum sequence length cutoff of 512 residues during training, truncating sequences that exceeded this threshold. Despite this limitation, the cutoff retains 172M (92%) of the annotated residues across the train, validation, and test sets, ensuring broad coverage of the dataset. Following (33, 71), we deploy LoRA (32) fine-tuning of the pLM during training with parameters *rank* = 4, *α* = 1 applied to query (Q), key (K), value (V), and output (O) layers using the Parameter-Efficient Fine-Tuning (PEFT) (33, 72) framework.

The supervised head consists of a lightweight convolutional neural network (CNN) with two hidden layers and two output heads: one for regression and one for classification. The first hidden layer (size = 64, kernel size = 7, padding = 3) is shared between both heads, while the second layer (size = 10, kernel size = 7, padding = 3) is task-specific, allowing the model to learn shared representations while capturing task-specific nuances. Each layer comprises a linear transformation, ReLU activation, and dropout with a probability of 0.1. The regression head predicts FrustratometeR-derived continuous frustration indices (FI) using mean squared error (MSE) loss. To mitigate class imbalance, the classification head of FrustrAI-Seq incorporates class-weighted cross-entropy (CE) loss to predict categorical frustration classes. Weights were derived from empirical class frequencies (73) and computed using the `compute_class_weight` function from scikit-learn. They are applied multiplicatively to the corresponding loss terms during training. The total loss is the sum of both individual losses, enabling joint optimization. To ensure accurate supervision, residues without frustration annotations are excluded from the loss computation during training.

We used the AdamW (74) optimizer with standard parameters (*β*_1_ = 0.9, *β*_2_ = 0.999) and an initial learning rate of 1e-4 with a warm-up over the first 500 steps. After that, we used cosine annealing to reduce the learning rate over the course of training to about 9e-5. All training was conducted on 4 Nvidia H100 GPUs, each with 80GB of memory, in distributed data parallel (DDP) mode, until convergence, with early stopping defined by a stagnating validation set loss over 5 validations. LoRA matrices comprise 2M parameters, approximately 0.16% of the total ProtT5 parameters. The supervised head has 3.2M parameters.

### 5.3. Baselines

The *random* baseline predictions were generated by independently shuffling the frustration scores and class labels and evaluating the resulting predictions against the true labels. Our second baseline is an amino acid-specific heuristic. For each residue *r*_*i*_ of type *a*, the predicted continuous frustration score is set to the mean frustration, *µ*_*a*_, observed for that amino acid *a* in the training set: 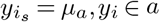. For the categorical frustration classes, each residue *x*_*i*_ is assigned the majority class 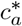 most frequently observed for amino acid 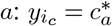.

### 5.4. Evalutation on Test Set

During training, FrustrAI-Seq is trained on the Funstration dataset, and a sequence cutoff is applied at a maximum sequence length of 512 (see Sec. 5.2). Its data loader indexes each residue by its position in the full UniProt full protein sequence and discards residues with `res_idx` ≥ 512 plus a small number lost to the ProtT5 end-of-sequence token at the boundary. Consequently FrustrAI-Seq scores 9,736,349 of the 10,654,715 test residues. The remaining 917,006 residues with `res_idx` ≥ 512 are outside its scope. We additionally ran an evaluation on the 917,006 residues only.

### 5.5. FrustraMPNN evaluation on the Funstration test set

We evaluated FrustraMPNN, a structure-based predictor of residue local energetic frustration built on top of the ProteinMPNN (36) message-passing encoder. We used the MegaScale checkpoint of FrustraMPNN and used our Funstration test set to evaluate performance. To report FrustraMPNN’s performance, we used the same 9,736,349 of the 10,654,715 test residues as in Sec5.4. Predicted indices were binned into three classes with the canonical thresholds: minimally frustrated (≥ 0.55), highly frustrated (≤−1.0), and neutral otherwise to compute F1, precision, and recall metrics.

### 5.6. Use cases inference

We evaluated two protein families: *α*-globins and *β*-lactamases. The *α*-globin family dataset was obtained from (9). Reference frustration-conservation results were generated using FrustraEvo (40) with a sequence distance (SeqDist) of 3, using PDB ID: 2DN1 (chain A) as the reference structure. A single reference structure was required to define the residues or contacts over which conservation was evaluated. Twelve protein–protein interaction (PPI) sites previously identified as highly frustrated at the family level were selected for mutational scanning. For each site, the native amino acid and all 19 non-native substitutions were evaluated, resulting in 240 residue-level cases (12 native and 228 mutant). For the *β*-lactamase use case, six characterized catalytic sites in the reference structure (PDB ID 4BLM, chain A) were selected. Similarly, the native amino acid and all 19 non-native substitutions were evaluated at each site, resulting in 120 residue-level cases (6 native and 114 mutant). Frustration changes for the *α*-globin and *β*-lactamase mutants were computed using FrustratometeR (11, 12) at SeqDist 3. Continuous frustration scores were inferred at the per-residue level for both the *α*-globin (2DN1) and the *β*-lactamase (4BLM) mutants using full-precision (32) on a single Nvidia H100 GPU.

### 5.7. De novo dataset

To create the de novo dataset, we used the Protein Design Archive keyword search (38) to identify four subsets of de novo-designed proteins: “enzyme”, “fibre”, “helical bundle”, and “nanoparticle” (e.g. “de novo AND enzyme”). All entries were downloaded on 22 June 2026. Further, all entries were split into single chains, duplicates were removed, and chains related to DNA were additionally removed, as FrustratometeR can not handle these. In the case of proteins that were part of two sets, they were removed from the helical bundle subset, as all of them were found in this set as well as in another, yielding 225 designs in total: 86 enzymes, 23 fibres, 102 helical bundles, and 14 nanoparticles. An extensive list of protein IDs is available in Supplementary Table 3.

### 5.8. ProteinMPNN designs

We re-used here Protein-MPNN sequences (36) designed already in prior work (39) for *α*-globins (n=21 experimental wildtype structures) and *β*-lactamases (n=31). Briefly, 20 sequences were generated for each native structure using ProteinMPNN T=0.1, and selected by higher ESMFold2 pLDDT. The same ESMFold structures were used to derive FrustratometeR frustration (using SeqDist3, no electrostatics).

### 5.9. Conformer dataset

For creating the conformer dataset, we searched the PDB (https://www.rcsb.org/) and PED (75) databases for protein ensembles containing more than 10 conformers, minimum length of 50 residues and spanning the full range of disorder content. To increase dataset diversity, particularly at high disorder content, we also included conformational ensembles generated by Jeon et al. (76). After merging the three sources, we clustered proteins at 80% sequence identity and used only representatives for further analysis. Disordered regions were predicted from protein sequences using SETH (26), retaining disorder annotation only for segments of at least 10 consecutive disordered residues (otherwise is defined as ordered). Protein disorder content was defined as the fraction of residues annotated as disordered. Ensembles were grouped into ten disorder-content bins, from 0–10% to 90–100%, keeping at most 50 ensembles per bin. The final dataset comprised 414 protein ensembles and 9,298 conformers.

### 5.10. Proteome-wide inference

Twenty-six reference proteomes were selected to provide broad taxonomic coverage. Proteomes represented in the AlphaFold Protein Structure Database (AFDB) were used as the starting set and supplemented with additional proteomes to improve coverage across taxonomic groups. The corresponding sequences were retrieved from Uniprot (69). Sequences containing non-canonical amino acids or exceeding 10,000 residues in length were excluded. For computational efficiency and performance reporting, continuous frustration scores and discrete frustration states were inferred at the per-residue level for all remaining sequences using half-precision (fp16) on a single Nvidia H100 GPU. The reported runtime denotes exclusively the inference time required by the model for a given filtered multifasta input. While the full proteomes UP000000554-64091, UP000001037-694429, UP000005640-9606, UP000006671-5762, UP000189677-193462, and UP000444721-5763 were processed individually, the remaining proteomes were collapsed and then splitted in bins of increasing protein length.

Intrinsic disorder was predicted with SETH (77) using default parameters, resulting in predicted values where 1 signifies maximum disorder and 0 maximum order. A threshold of 0.5 was chosen to separate residues into ordered (≤ 0.5) and disordered (> 0.5).

In a separate large-scale analysis, 22,105 reference proteomes, comprising 136,775,502 proteins, were retrieved from Uniprot. Per-residue frustration was predicted with FrustrAI-Seq on a single Nvidia H100 GPU, app.ying the same sequecne-filtering criteria and maximum length of 10,000 residues. For each protein, the fraction of residues classified as highly, neutral or minimally frstarades were caluclated normalizing by protein length. Proteins were ranked independently according to each fraction, and the top 1 % of each distribution was retained. Because these sets remained too large for detailed analysis, 5,000 proteins were sampled evenly across the ranked top 1 % of each frustration category for the supplementary analysis. Independently, the complete dataset was filtered to retain proteins supported by evidence at the protein or transcript level, reducing it from approximately 136 million to 255,000 proteins while spanning the same 22,105 proteomes. For the runtime analysis, FrustratometeR was run using a single core of an Intel(R) Xeon(R) Platinum 8480+ CPU.

### 5.11. Post-translational modifications

For the post-translational modification (PTM) analysis, CATH domains in the Funstration dataset were mapped to their corresponding UniProt sequences and PTM annotations for phosphorilation and glycosylation sites were transferred at residue level based on matching residue position and amino-acid identity. Residues were divided into PTM-annotated and unannotated sets. To control for differences in amino-acid composition, residues from the unannotated set were randomly sampled to match the sample size and amino-acid distribution of the PTM-annotated set. The proportions of highly frustrated, neutral, and minimally frustrated residues were then calculated for both sets using the FrustratometeR frustration index and the FrustrAI-Seq regression predictions.

### 5.12. Metrics

We calculated Precision, Recall, and the F1 score to evaluate the classification head’s performance. They are defined as

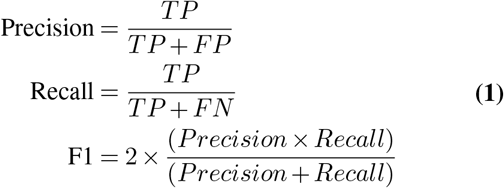

where TP are True Positives, FP are False Positives, and FN are False Negatives. Macro scores are the unweighted averages across all classes.

To estimate reliabilities, we use Shannon Entropy on the softmax-transformed outputs of the classification heads’ last layer output logits *ŵ*.

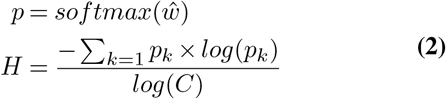

where *p*_*k*_ denotes the predicted probability of class *k*, and *C* denotes the number of classes, which is 3 in our case.

The discovery score *z* is an amino acid type-specific z-score transformation.

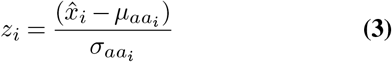

where *aa*_*i*_ ∈ *AA* denotes the amino acid type of residue *i*, and 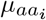 and 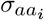 are the mean and standard deviation for that amino acid class. We computed *µ*_*aa*_ and *σ*_*aa*_ on the training set for each amino acid.

To quantify the agreement between predicted and ground-truth frustration profiles at the protein level, we computed the cosine similarity between the residue-wise Frustration Index (FI) vectors. For a protein of length *L*, let 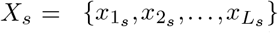 denote the FrustratometeR-derived frustration values, and 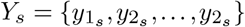 the corresponding FrustrAI-Seq predictions. The cosine similarity is defined as

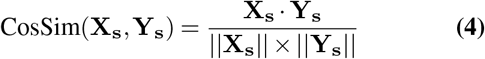

To assess residue-level disagreement between predicted and ground-truth frustration classes, we computed the Hamming distance. Let 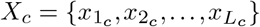 denote the ground-truth and 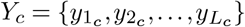 predicted class labels, respectively. The Hamming distance is defined as

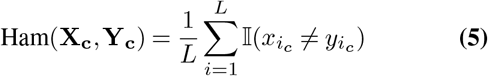

where I(·) is the indicator function. Both Cosine similarity and Hamming distance were computed per protein over the test set.

## Competing interests

No competing interest is declared.

## Author contributions statement

RGP and MH conceived the experiments; NB and RGP prepared the dataset; JPL trained the models; JPL, MPG, MF, JS, and FLS conducted analyses; all authors wrote the manuscript.

## Acknowledgments

This work was supported by Helmholtz AI computing resources (HAICORE) of the Helmholtz Association’s Initiative and Networking Fund through Helmholtz AI. RGP is funded by the Ramon y Cajal program (RYC2023-043825-I) and the MEGAFrustratEDS grant (PID2024-159128OA-I00). MPG is funded by a COLLABORATION CONTRACT FOR RESEARCH AND DEVELOPMENT SERVICES IN HPC signed between the Barcelona Supercomputing Center and NVIDIA Corporation on 15/12/2023. MFM was supported by Grant PRE2022-101718 funded by MICIU/AEI/10.13039/501100011033 and by “ESF+” Grant CEX2021-001148-S-20-2 funded by MI-CIU/AEI/10.13039/501100011033. FLS is a fellow within the “Generación D” initiative, Red.es, Ministerio para la Transformación Digital y de la Función Pública, for talent attraction (C005/24-ED CV1). Funded by the European Union NextGenerationEU funds, through PRTR.

## Notes

https://github.com/leuschjanphilipp/FrustrAI-Seq

### Competing Interest Statement

The authors have declared no competing interest.

### Summary of Updates

This is a revised version of the initial manuscript, including several new insights, such as a coformer analysis, frustration-disorder coupling, and large-scale proteome analysis, along with a revised discussion. The author list has been extended.

