## Supplementary material for "FrustrAI-Seq: Scaling Local Energetic Frustration to the Protein Sequence Space": FrustrAI-Seq_Supplementary

DRAFT

**Supplementary Table 1. Ablation study.** We monitored the effect of various design choices of the final FrustrAI-Seq using F1-score, precision, recall, Pearson's $r$, and mean-absolute error (MAE) on our validation set. The top block reports metrics derived from the *binned regression head*; the bottom block reports metrics derived from the *classification head*. We established a) ProtT5 outperforms ESM2-650M and ProstT5 numerically, b) having combined classification (CLS) and regression (REG) loss (as in row *ProtT5*) performs better than having either of them, c) adding class-weights (CW) to the cross-entropy loss for highly-frustrated residues trades precision for recall, d) adding low-ranked adapters (LoRA) generally improves performance, and e), using Focal-loss instead of class-weights has a similar effect but reaches slightly lower recall for highly frustrated residues. Taken together, our final model FrustrAI-Seq LoRA-finetunes ProtT5 using simultaneously a regression and classification head with class-weights applied to the cross-entropy loss to improve recall of the minority class. Pearson $r$ and MAE are omitted from the classification block as they are always based on the regression heads' output.

| | Macro | | | Highly Frust | | | Neutral | | | Minimally Frust | | | Pear. | |
|---|---|---|---|---|---|---|---|---|---|---|---|---|---|---|
| | F1 | Prec. | Rec. | F1 | Prec. | Rec. | F1 | Prec. | Rec. | F1 | Prec. | Rec. | $r$ | MAE |
| *Regression Head* | | | | | | | | | | | | | | |
| ESM2 650M | 0.68 | 0.78 | 0.65 | 0.40 | 0.71 | 0.28 | 0.80 | 0.70 | 0.93 | 0.82 | 0.93 | 0.74 | 0.83 | 0.38 |
| ProstT5 | 0.65 | 0.76 | 0.63 | 0.33 | 0.66 | 0.22 | 0.79 | 0.69 | 0.93 | 0.82 | 0.93 | 0.74 | 0.81 | 0.40 |
| ProtT5 | 0.70 | 0.79 | 0.67 | 0.44 | 0.74 | 0.32 | 0.81 | 0.72 | 0.93 | 0.84 | 0.93 | 0.77 | 0.84 | 0.36 |
| ProtT5, CLS only | – | – | – | – | – | – | – | – | – | – | – | – | – | – |
| ProtT5, REG only | 0.71 | 0.80 | 0.68 | 0.47 | 0.75 | 0.34 | 0.81 | 0.73 | **0.91** | 0.85 | 0.91 | 0.79 | 0.85 | 0.35 |
| ProtT5, w CW | 0.69 | 0.80 | 0.67 | 0.44 | 0.75 | 0.31 | 0.81 | 0.72 | 0.93 | 0.84 | **0.93** | 0.77 | 0.85 | 0.36 |
| ProtT5, LoRA, w/o CW | **0.74** | **0.81** | **0.72** | **0.55** | **0.75** | **0.44** | 0.82 | 0.75 | 0.91 | 0.86 | 0.92 | 0.81 | 0.87 | 0.33 |
| ProtT5, LoRA, Focal loss | 0.74 | 0.81 | 0.72 | 0.55 | 0.75 | 0.44 | **0.82** | 0.75 | 0.90 | **0.86** | 0.91 | **0.81** | **0.87** | **0.32** |
| **FrustrAI-Seq** | 0.74 | 0.81 | 0.71 | 0.55 | 0.75 | 0.43 | 0.82 | 0.75 | 0.90 | 0.86 | 0.91 | 0.81 | 0.86 | 0.33 |
| *Classification Head* | | | | | | | | | | | | | | |
| ESM2 650M | 0.73 | 0.76 | 0.71 | 0.54 | 0.66 | 0.46 | 0.80 | 0.75 | 0.86 | 0.84 | 0.88 | 0.81 | – | – |
| ProstT5 | 0.70 | 0.75 | 0.68 | 0.48 | 0.64 | 0.38 | 0.79 | 0.73 | 0.88 | 0.84 | 0.89 | 0.79 | – | – |
| ProtT5 | 0.74 | 0.77 | 0.73 | 0.57 | 0.65 | 0.50 | 0.81 | 0.77 | 0.85 | 0.86 | 0.88 | 0.83 | – | – |
| ProtT5, CLS only | 0.74 | 0.78 | 0.72 | 0.55 | 0.68 | 0.46 | 0.81 | 0.76 | 0.87 | 0.85 | 0.89 | 0.82 | – | – |
| ProtT5, REG only | 0.71 | 0.80 | 0.68 | 0.47 | **0.75** | 0.34 | 0.81 | 0.73 | **0.91** | 0.85 | **0.91** | 0.79 | – | – |
| ProtT5, w CW | 0.73 | 0.72 | 0.78 | 0.59 | 0.45 | **0.85** | 0.74 | 0.85 | 0.65 | 0.86 | 0.86 | 0.85 | – | – |
| ProtT5, LoRA, w/o CW | **0.77** | **0.79** | 0.76 | **0.63** | 0.68 | 0.59 | **0.82** | 0.79 | 0.85 | **0.87** | 0.89 | 0.85 | – | – |
| ProtT5, LoRA, Focal loss | 0.75 | 0.73 | 0.79 | 0.62 | 0.49 | 0.83 | 0.75 | 0.84 | 0.68 | 0.86 | 0.86 | **0.87** | – | – |
| **FrustrAI-Seq** | 0.75 | 0.74 | **0.80** | 0.62 | 0.49 | 0.85 | 0.77 | **0.85** | 0.69 | 0.87 | 0.87 | 0.86 | – | – |

**Supplementary Table 2. De novo evaluation.** We report F1-score, precision, recall, Pearson's $r$, and mean absolute error (MAE) for FrustrAI-Seq on de novo datasets from the Protein Design Archive, compared against the Frustratometer reference. The top block reports metrics from the *regression head*; the bottom block reports metrics from the *classification head*. The first row of each block corresponds to performance over all de novo sets. Pearson $r$ and MAE are omitted from the classification block as they are always based on the regression heads' output.

| | Macro | | | Highly Frust | | | Neutral | | | Minimally Frust | | | Pear. | |
|---|---|---|---|---|---|---|---|---|---|---|---|---|---|---|
| | F1 | Prec. | Rec. | F1 | Prec. | Rec. | F1 | Prec. | Rec. | F1 | Prec. | Rec. | $r$ | MAE |
| *Regression Head* | | | | | | | | | | | | | | |
| All de novo | 0.72 | 0.77 | 0.69 | 0.52 | 0.71 | 0.40 | 0.78 | 0.71 | 0.86 | 0.86 | 0.90 | 0.82 | 0.87 | 0.33 |
| Enzyme | 0.71 | 0.78 | 0.69 | 0.49 | 0.70 | 0.37 | 0.80 | 0.74 | 0.88 | 0.85 | 0.89 | 0.82 | 0.86 | 0.34 |
| Fibre | 0.74 | 0.79 | 0.72 | 0.57 | 0.75 | 0.46 | 0.78 | 0.70 | 0.88 | 0.88 | 0.93 | 0.83 | 0.88 | 0.32 |
| Helical bundle | 0.71 | 0.76 | 0.69 | 0.52 | 0.71 | 0.41 | 0.76 | 0.68 | 0.85 | 0.85 | 0.90 | 0.81 | 0.88 | 0.32 |
| Nanoparticle | 0.73 | 0.77 | 0.71 | 0.54 | 0.70 | 0.44 | 0.76 | 0.69 | 0.85 | 0.87 | 0.91 | 0.84 | 0.89 | 0.30 |
| *Classification Head* | | | | | | | | | | | | | | |
| All de novo | 0.72 | 0.71 | 0.77 | 0.60 | 0.48 | 0.82 | 0.69 | 0.80 | 0.61 | 0.87 | 0.86 | 0.88 | – | – |
| Enzyme | 0.72 | 0.71 | 0.77 | 0.57 | 0.44 | 0.79 | 0.73 | 0.82 | 0.66 | 0.87 | 0.86 | 0.87 | – | – |
| Fibre | 0.74 | 0.73 | 0.79 | 0.65 | 0.52 | 0.84 | 0.69 | 0.79 | 0.62 | 0.89 | 0.89 | 0.89 | – | – |
| Helical bundle | 0.71 | 0.71 | 0.76 | 0.61 | 0.48 | 0.84 | 0.66 | 0.78 | 0.57 | 0.87 | 0.85 | 0.88 | – | – |
| Nanoparticle | 0.74 | 0.73 | 0.78 | 0.64 | 0.51 | 0.86 | 0.68 | 0.80 | 0.60 | 0.89 | 0.88 | 0.90 | – | – |

**Supplementary Table 3. De novo dataset composition.** PDB identifiers of the de novo designed proteins used for evaluation, grouped by structural class as annotated in the Protein Design Archive (enzymes, fibres, helical bundles, and nanoparticles). Each identifier is given as `<PDB ID>_<chain>`. In total the de novo set comprises 225 chains (86 enzymes, 23 fibres, 102 helical bundles, and 14 nanoparticles).

| Class | PDB identifiers |
|---|---|
| Enzyme (86) | 2kl8_A, 2lze_A, 2n2t_A, 2n76_A, 3pbj_A, 3pbj_B, 3u0s_A, 4ky3_A, 4ky3_B, 4kyz_A, 4yxx_A, 4yxy_A, 4yxz_A, 4yy2_C, 4yy5_A, 4yy5_B, 5bvl_A, 5byo_A, 5cw9_A, 5vjs_A, 5vju_A, 6c2v_A, 6nuk_A, 6nuk_B, 6wi5_A, 6wxo_A, 6wxo_B, 6yqx_A, 6yqy_A, 7a8s_A, 7l33_A, 7mcc_A, 7mcd_A, 7mcd_B, 7mcd_C, 7mwq_A, 7mwq_B, 7osu_A, 7p12_A, 7smj_A, 7uek_A, 8c3w_A, 8dt0_A, 8dt0_B, 8fje_A, 8fjf_A, 8fjg_A, 8fre_A, 8frf_A, 8frf_C, 8frf_D, 8frf_E, 8frf_F, 8frf_G, 8frf_H, 8k7m_A, 8k7m_C, 8kc8_A, 8kdq_A, 8oys_A, 8tcf_A, 8tcf_B, 8tcf_C, 8vhs_A, 8vhs_D, 9bqr_A, 9fw5_A, 9fw7_A, 9fwa_A, 9gbt_A, 9gvf_A, 9jix_A, 9jix_C, 9mrb_A, 9pyj_A, 9pyj_B, 9pyj_D, 9pyl_A, 9pyl_B, 9qdp_A, 9r00_A, 9r7f_A, 9szu_A, 9ugr_A, 9ukw_A, 9ytq_A |
| Fibre (23) | 3ra3_A, 3ra3_B, 3ra3_D, 3ra3_E, 8eov_A, 8eox_A, 8eox_B, 8eox_D, 8eoz_B, 8erw_A, 8erw_B, 8g8i_A, 8gaa_A, 8gaq_A, 8uao_A, 8ub3_A, 8ubg_A, 9cc5_A, 9cc5_B, 9cc6_A, 9cc6_B, 9nzh_A, 9nzh_B |
| Helical bundle (102) | 12td_A, 1byz_A, 2lse_A, 4tql_A, 4tql_B, 4uos_A, 4uot_A, 5cwi_A, 5cwm_A, 5wlj_A, 5wlk_A, 5wll_A, 5wlm_A, 6e9t_B, 6e9v_Q, 6e9z_B, 6egc_A, 6g65_A, 6g66_A, 6g67_A, 6g68_A, 6g68_G, 6g69_A, 6g69_H, 6g69_N, 6g6a_A, 6g6a_C, 6g6b_A, 6g6b_B, 6g6c_A, 6g6c_G, 6g6c_J, 6g6d_A, 6g6e_A, 6g6f_A, 6g6f_D, 6g6g_A, 6g6h_A, 6mq2_A, 6mq2_D, 6n4n_A, 6n4n_C, 6n4n_F, 6u1s_A, 6xi6_A, 6xr2_A, 6xr2_B, 6xr2_C, 6xr2_D, 6xr2_E, 6xr2_F, 6xt4_A, 7cbc_A, 7cbc_B, 7jh5_A, 7jh5_B, 7udj_G, 7udj_H, 7udo_A, 7unh_A, 7unh_B, 7uni_A, 7uni_B, 7uni_C, 7uni_D, 7unj_A, 7unj_B, 8e0l_A, 8e0l_C, 8e0o_A, 8e0o_B, 8e0o_C, 8f6q_A, 8f6r_A, 8flx_A, 8ga6_A, 8gel_A, 8gl3_A, 8qae_A, 8qae_C, 8qag_A, 8qag_B, 8qkd_A, 8tl7_A, 8tnb_A, 8u1a_A, 8u1a_B, 8ugc_A, 8utm_A, 8v2d_J, 8v2d_y, 8v3b_a, 8vej_A, 8vej_B, 8vx7_A, 8vx7_B, 9dd1_C, 9dt9_A, 9exk_A, 9hnh_A, 9nlt_A, 9nlt_B |
| Nanoparticle (14) | 3s0r_A, 4qtr_A, 4qtr_B, 4qtr_C, 8cyk_A, 8cyk_B, 8f4x_0, 8f54_L, 8fbi_A, 8fbj_A, 9dze_K, 9dze_a, 9dze_f, 9ndl_A |

**Supplementary Table 4. Bootstrap uncertainty of the binned-regression-head performance metrics (main Table 1).** For every metric we report the point estimate together with its bootstrap standard error (SE) as *value* ± *SE*. The discrete frustration class is obtained by thresholding the continuous regression output at FI $= -1$ and $0.55$ (class 0 = highly, 1 = neutral, 2 = minimally frustrated); the point estimates reproduce the corresponding entries of Table 1 in the main text. Uncertainties were obtained by a grouped, protein-level bootstrap of the hold-out test set (1000 replicates, whole proteins resampled with replacement; see the Metrics section of the main-text Methods). Macro F1, precision and recall are the unweighted means over the three frustration classes; per-class values are added for completeness. Models are ordered as in Table 1; the final model *FrustrAI-Seq* is highlighted in bold. †FrustraMPNN predicts frustration from 3D backbones. Every standard error is below $0.001$, more than an order of magnitude smaller than the performance differences between models, so all comparisons discussed in the text are statistically unambiguous.

| | FrustraMPNN† | ESM2 650M | ProstT5 | ProtT5 | **FrustrAI-Seq** |
|---|---|---|---|---|---|
| *Macro average* | | | | | |
| F1 | $0.677 \pm 0.0002$ | $0.681 \pm 0.0003$ | $0.650 \pm 0.0002$ | $0.699 \pm 0.0003$ | $\mathbf{0.739 \pm 0.0003}$ |
| Prec. | $0.689 \pm 0.0002$ | $0.783 \pm 0.0003$ | $0.762 \pm 0.0003$ | $0.797 \pm 0.0002$ | $\mathbf{0.807 \pm 0.0002}$ |
| Rec. | $0.669 \pm 0.0002$ | $0.654 \pm 0.0003$ | $0.628 \pm 0.0002$ | $0.670 \pm 0.0003$ | $\mathbf{0.710 \pm 0.0003}$ |
| *Highly frustrated* | | | | | |
| F1 | $0.454 \pm 0.0005$ | $0.420 \pm 0.0007$ | $0.337 \pm 0.0006$ | $0.449 \pm 0.0007$ | $\mathbf{0.542 \pm 0.0006}$ |
| Prec. | $0.495 \pm 0.0006$ | $0.711 \pm 0.0007$ | $0.669 \pm 0.0009$ | $0.747 \pm 0.0007$ | $\mathbf{0.762 \pm 0.0005}$ |
| Rec. | $0.420 \pm 0.0005$ | $0.298 \pm 0.0006$ | $0.225 \pm 0.0005$ | $0.321 \pm 0.0006$ | $\mathbf{0.420 \pm 0.0007}$ |
| *Neutral* | | | | | |
| F1 | $0.764 \pm 0.0002$ | $0.802 \pm 0.0001$ | $0.794 \pm 0.0001$ | $0.811 \pm 0.0001$ | $\mathbf{0.822 \pm 0.0001}$ |
| Prec. | $0.735 \pm 0.0002$ | $0.706 \pm 0.0002$ | $0.695 \pm 0.0002$ | $0.721 \pm 0.0002$ | $\mathbf{0.752 \pm 0.0002}$ |
| Rec. | $0.796 \pm 0.0002$ | $0.929 \pm 0.0001$ | $0.925 \pm 0.0001$ | $0.925 \pm 0.0001$ | $\mathbf{0.906 \pm 0.0002}$ |
| *Minimally frustrated* | | | | | |
| F1 | $0.813 \pm 0.0002$ | $0.822 \pm 0.0002$ | $0.818 \pm 0.0002$ | $0.837 \pm 0.0002$ | $\mathbf{0.853 \pm 0.0002}$ |
| Prec. | $0.837 \pm 0.0002$ | $0.932 \pm 0.0002$ | $0.922 \pm 0.0002$ | $0.924 \pm 0.0002$ | $\mathbf{0.908 \pm 0.0002}$ |
| Rec. | $0.790 \pm 0.0003$ | $0.735 \pm 0.0003$ | $0.735 \pm 0.0003$ | $0.765 \pm 0.0003$ | $\mathbf{0.805 \pm 0.0003}$ |
| *Regression head* | | | | | |
| Pearson $r$ | $0.764 \pm 0.0002$ | $0.824 \pm 0.0002$ | $0.806 \pm 0.0002$ | $0.839 \pm 0.0002$ | $\mathbf{0.859 \pm 0.0002}$ |
| MAE | $0.462 \pm 0.0002$ | $0.388 \pm 0.0002$ | $0.408 \pm 0.0002$ | $0.366 \pm 0.0002$ | $\mathbf{0.336 \pm 0.0002}$ |

**Supplementary Table 5. Bootstrap uncertainty of the classification-head performance metrics.** For every metric we report the point estimate together with its bootstrap standard error (SE) as *value* ± *SE*. Uncertainties were obtained by a grouped, protein-level bootstrap of the hold-out test set (1000 replicates, whole proteins resampled with replacement; see the Metrics section of the main-text Methods). Macro F1, precision and recall are the unweighted means over the three frustration classes. per-class values and the regression-head Pearson $r$ and MAE are added for completeness. Models are ordered as in Table 1 of the main text; the final model *FrustrAI-Seq* is highlighted in bold. †FrustraMPNN predicts frustration from 3D backbones. Every standard error is below $0.001$, more than an order of magnitude smaller than the performance differences between models, so all comparisons discussed in the text are statistically unambiguous.

| | FrustraMPNN† | ESM2 650M | ProstT5 | ProtT5 | **FrustrAI-Seq** |
|---|---|---|---|---|---|
| *Macro average* | | | | | |
| F1 | $0.677 \pm 0.0002$ | $0.728 \pm 0.0002$ | $0.700 \pm 0.0002$ | $0.742 \pm 0.0002$ | $\mathbf{0.753 \pm 0.0002}$ |
| Prec. | $0.689 \pm 0.0002$ | $0.761 \pm 0.0002$ | $0.750 \pm 0.0002$ | $0.769 \pm 0.0002$ | $\mathbf{0.738 \pm 0.0002}$ |
| Rec. | $0.669 \pm 0.0002$ | $0.709 \pm 0.0002$ | $0.678 \pm 0.0002$ | $0.725 \pm 0.0002$ | $\mathbf{0.802 \pm 0.0002}$ |
| *Highly frustrated* | | | | | |
| F1 | $0.454 \pm 0.0005$ | $0.539 \pm 0.0005$ | $0.472 \pm 0.0005$ | $0.563 \pm 0.0005$ | $\mathbf{0.623 \pm 0.0004}$ |
| Prec. | $0.495 \pm 0.0006$ | $0.653 \pm 0.0006$ | $0.632 \pm 0.0007$ | $0.659 \pm 0.0006$ | $\mathbf{0.495 \pm 0.0005}$ |
| Rec. | $0.420 \pm 0.0005$ | $0.459 \pm 0.0006$ | $0.377 \pm 0.0005$ | $0.491 \pm 0.0006$ | $\mathbf{0.842 \pm 0.0004}$ |
| *Neutral* | | | | | |
| F1 | $0.764 \pm 0.0002$ | $0.805 \pm 0.0002$ | $0.796 \pm 0.0002$ | $0.811 \pm 0.0002$ | $\mathbf{0.773 \pm 0.0002}$ |
| Prec. | $0.735 \pm 0.0002$ | $0.757 \pm 0.0002$ | $0.731 \pm 0.0002$ | $0.772 \pm 0.0002$ | $\mathbf{0.857 \pm 0.0002}$ |
| Rec. | $0.796 \pm 0.0002$ | $0.860 \pm 0.0002$ | $0.873 \pm 0.0002$ | $0.855 \pm 0.0002$ | $\mathbf{0.705 \pm 0.0002}$ |
| *Minimally frustrated* | | | | | |
| F1 | $0.813 \pm 0.0002$ | $0.839 \pm 0.0002$ | $0.832 \pm 0.0002$ | $0.852 \pm 0.0002$ | $\mathbf{0.861 \pm 0.0002}$ |
| Prec. | $0.837 \pm 0.0002$ | $0.873 \pm 0.0002$ | $0.886 \pm 0.0002$ | $0.876 \pm 0.0002$ | $\mathbf{0.863 \pm 0.0002}$ |
| Rec. | $0.790 \pm 0.0003$ | $0.807 \pm 0.0003$ | $0.784 \pm 0.0003$ | $0.829 \pm 0.0002$ | $\mathbf{0.860 \pm 0.0002}$ |

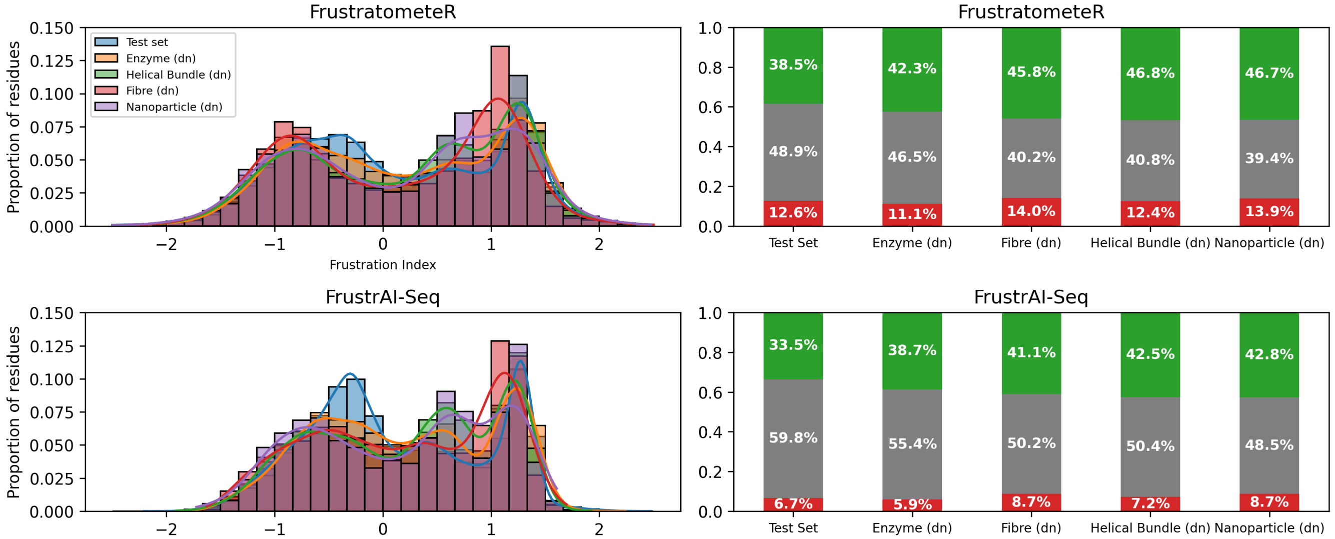


**Supplementary Figure 1. Frustration Index Distribution of FrustratometeR and FrustrAI-Seq across Funstration test set and de novo datasets.** The left histograms show the continuous FI score distributions for each dataset, normalized by the number of residues in each dataset for FrustratometeR (top) and FrustrAI-Seq (bottom). Right bar plots show the class attribution if the continuous scores are discretized into the three frustration states at thresholds $-1$ and $0.55$. Red indicates highly frustrated residues, gray indicates neutral residues, and green indicates minimally frustrated residues.

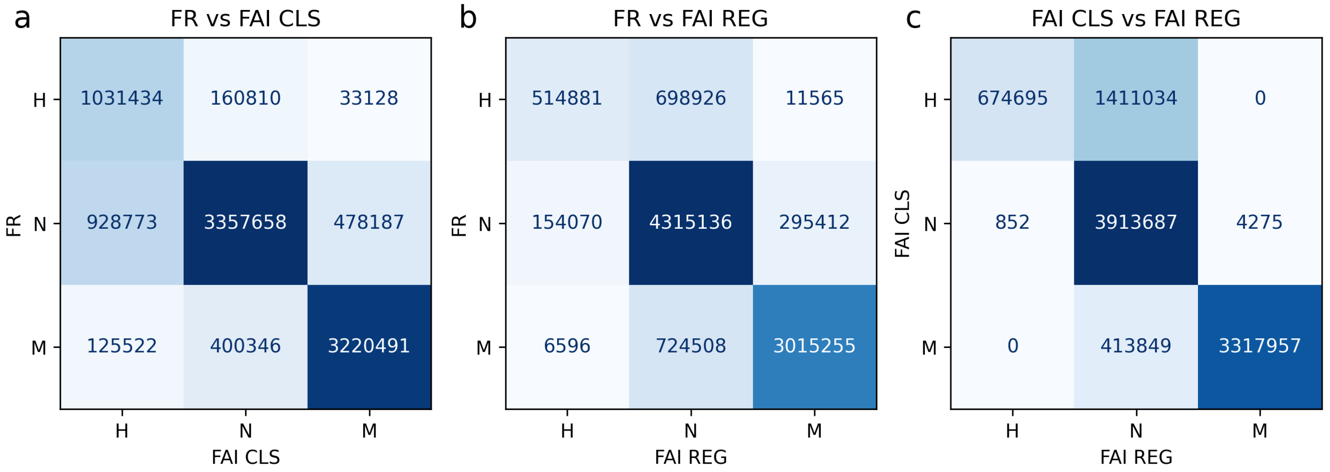


**Supplementary Figure 2. Comparison of FrustrAI-Seq's classification and regression head outputs to FrustratometeR.** FR corresponds to FrustratometeR outputs discretized with thresholds $-1$ and $0.55$. FAI CLS is the output of the FrustrAI-Seq classification head; FAI REG is FrustrAI-Seq's regression head output discretized following the same thresholds as FrustratometeR. The three classes H, N, and M represent the frustration states: highly frustrated, neutral, and minimally frustrated. (a) depicts a confusion matrix comparing FAI CLS head outputs to their test set labels from FrustratometeR. (b) shows the confusion matrix for FrustrAI-Seq's binned regression predictions against FrustratometeR. (c) compares the agreement between FrustrAI-Seq's regression and classification heads. All data are based on the Funstration test set, considering only residues below position 512 in their corresponding protein sequences, mirroring the approach used for training.

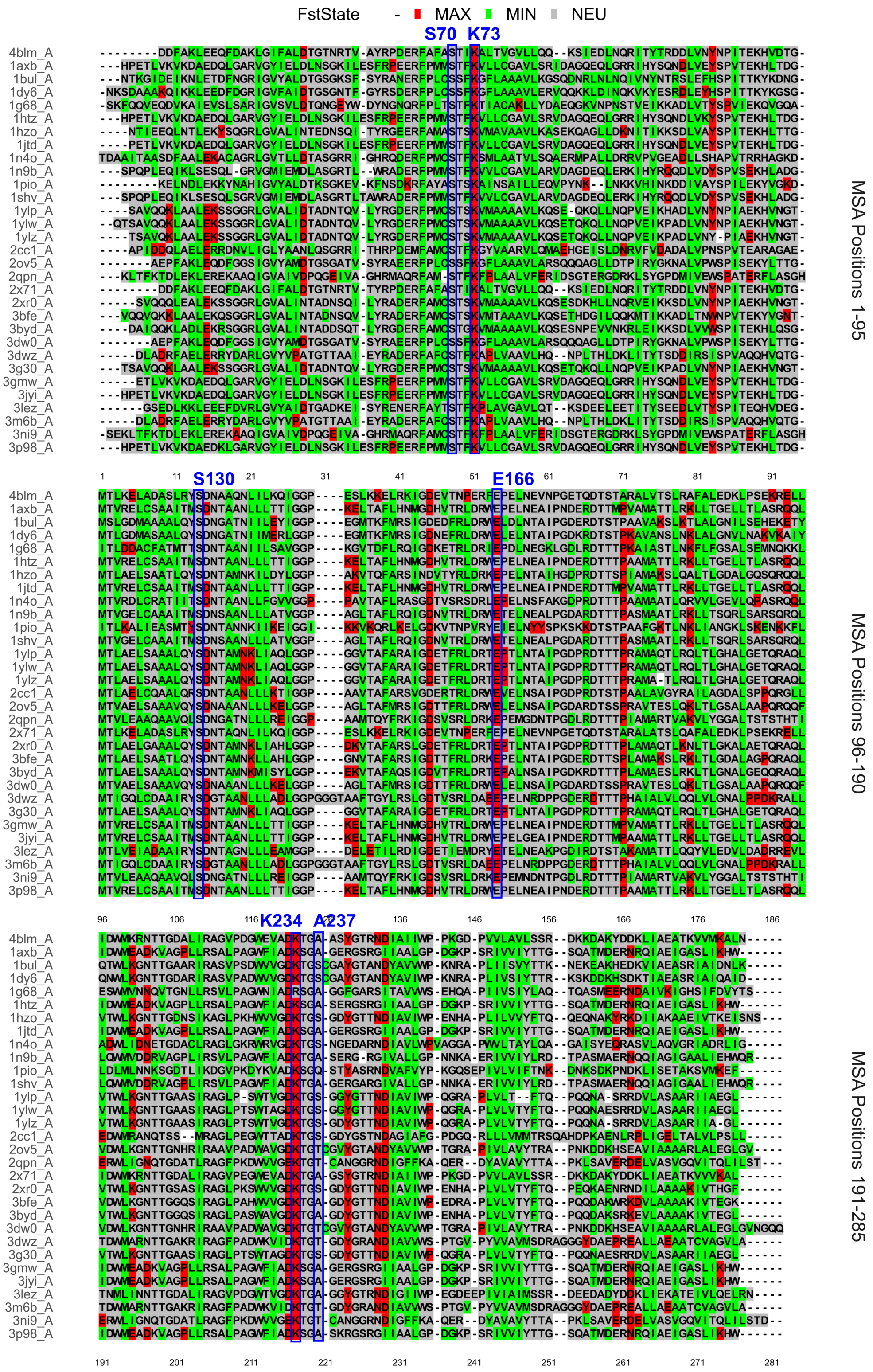


**Supplementary Figure 3. FrustrAI-Seq prediction of local frustration in $\beta$-Lactamases. Mutiple Sequence Frustration Alignment (MSFA):** Local frustration predictions by FrustrAI-seq projected on top of the $\beta$-Lactamases multiple sequence alignment. Columns corresponding to the catalytic residues are marked in blue rectangles

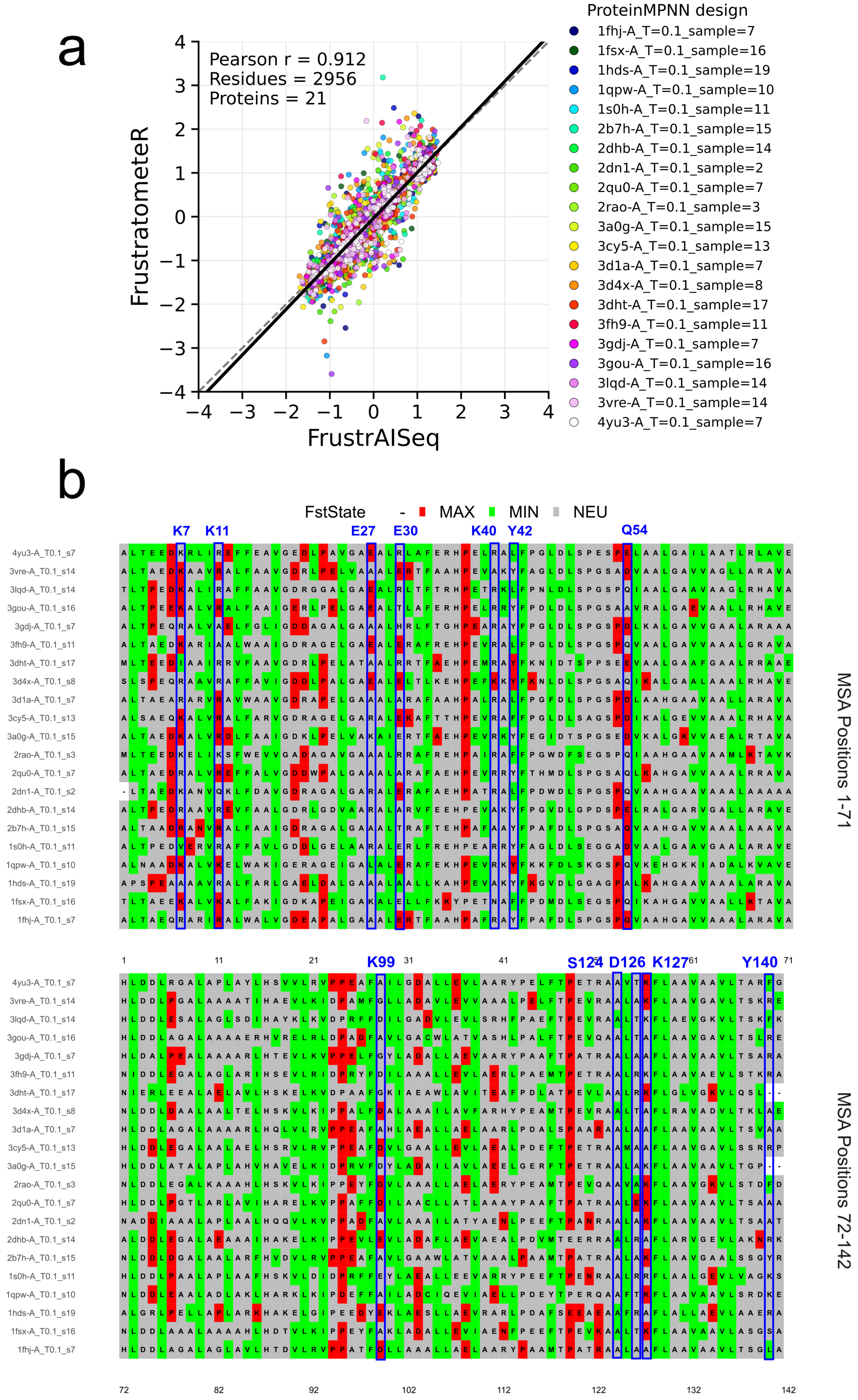


**Supplementary Figure 4. FrustrAI-Seq prediction of local frustration in reverse-folded $\alpha$-globins by ProteinMPNN.** (a) Scatter plot comparing the frustration index predicted by FrustrAI-Seq (x-axis) with the reference frustration index calculated by Frustratometer (y-axis) for all residues across the $\alpha$-globin synthetic family (n=21, as the native one). Points are colored by protein members, being here selected designs from ProteinMPNN runs. The Pearson correlation coefficient (r), along with the total number of residues and proteins analyzed, is displayed in the upper-left corner. (b) Mutiple Sequence Frustration Alignment (MSFA) mapping local frustration predictions by FrustrAI-seq on top of the $\alpha$-globin MSA containing ProteinMPNN designs instead of the native family. Frustration states are derived from standard threshold-based residue classification: green, minimally frustrated; red, highly frustrated; gray, neutral. Columns involved in PPIs (K7, K11, E27, E30, K40, Y42, Q54, K99, S124, D126, K127 and Y140) are marked in blue rectangles.

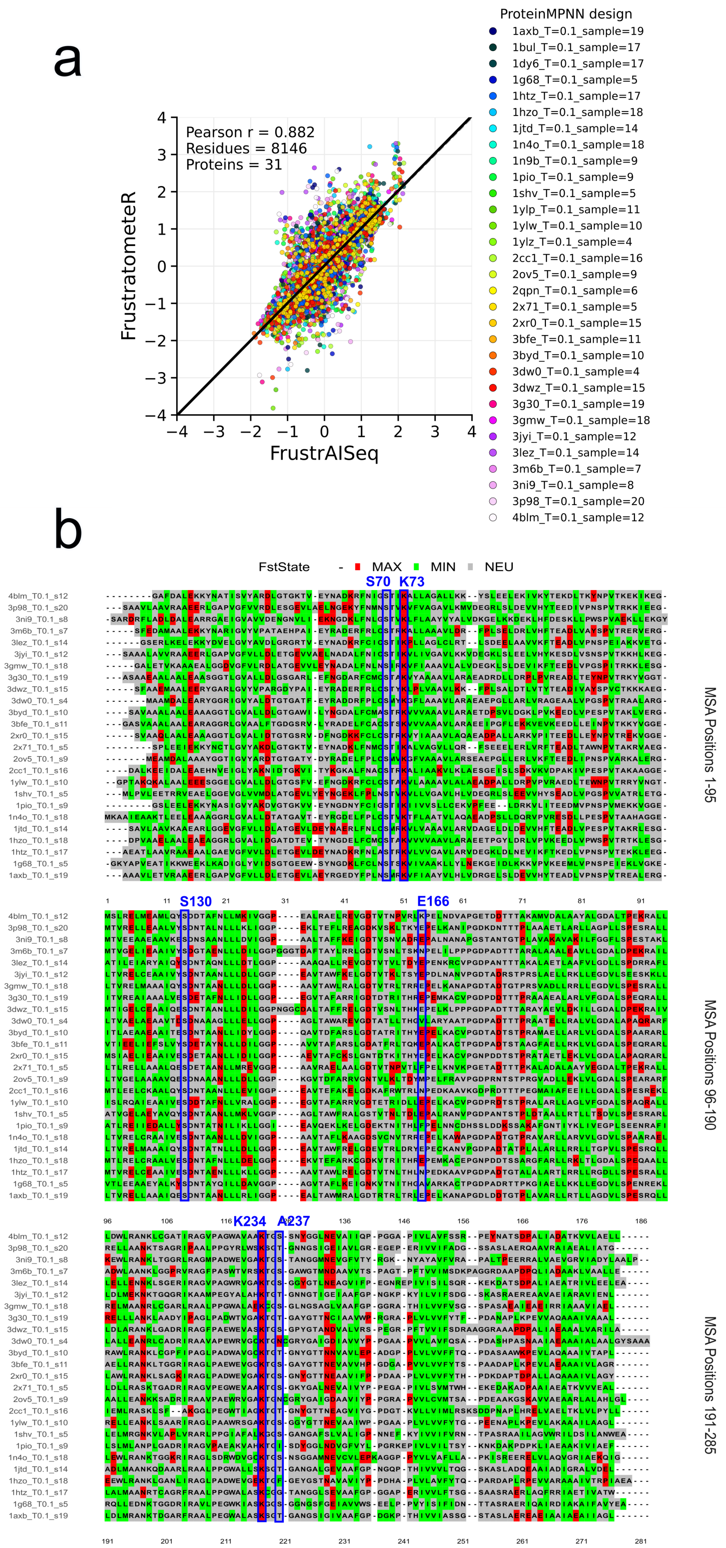


**Supplementary Figure 5. FrustrAI-Seq prediction of local frustration in reverse-folded $\beta$-Lactamases by ProteinMPNN.** (a) Scatter plot comparing the frustration index predicted by FrustrAI-Seq (x-axis) with the reference frustration index calculated by Frustratometer (y-axis) for all residues across the $\beta$-Lactamases synthetic family (n=31, as the native one). Points are colored by protein members, being here selected designs from ProteinMPNN runs. The Pearson correlation coefficient (r), along with the total number of residues and proteins analyzed, is displayed in the upper-left corner. (b) Mutiple Sequence Frustration Alignment (MSFA) mapping local frustration predictions by FrustrAI-seq on top of the $\beta$-Lactamases MSA containing ProteinMPNN designs instead of the native family. 6 out of 31 family members were excluded as ProteinMPNN redesigned more residues for them than were present in the native MSA used for the original FrustraEvo. This discrepancy caused structural and positional inconsistencies when constructing the template-based MSA. Frustration states are derived from standard threshold-based residue classification: green, minimally frustrated; red, highly frustrated; gray, neutral. Columns corresponding to the catalytic residues are marked in blue rectangles.

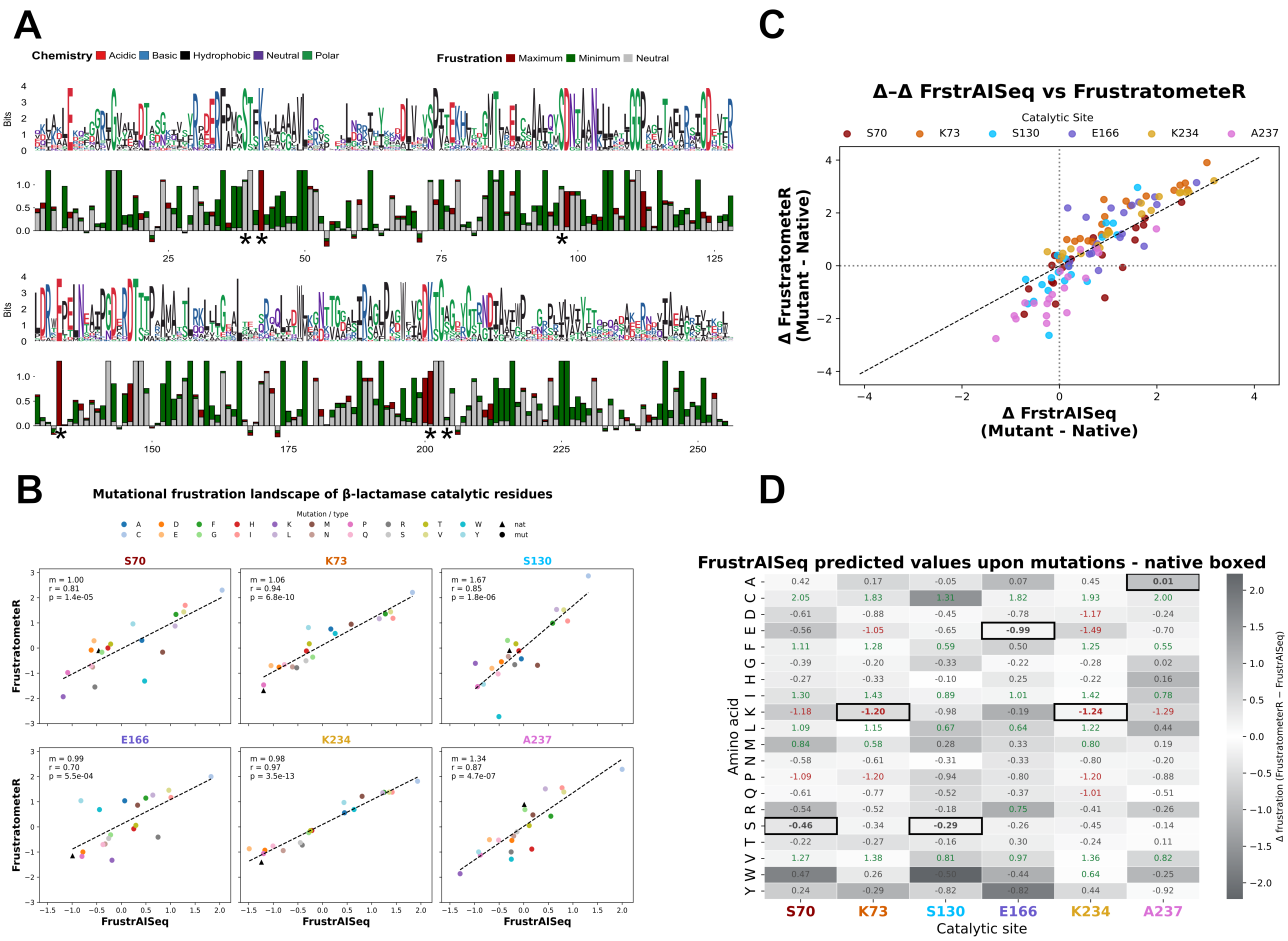


**Supplementary Figure 6. Sensitivity of FrustrAI-Seq to point mutations in $\beta$-lactamases. (A)** Frustration logo for the $\beta$-lactamase family, integrating sequence conservation (1st and 3rd row) and predicted frustration states (2nd and 4th row) across homologs. Conserved frustration patterns are observed at functionally and structurally constrained sites, corresponding to catalytic residues (positions 70, 73, 130, 166, 234, and 237 and marked with *). **(B)** Comparison of frustration values predicted by FrustrAI-Seq and computed by FrustratometeR for six catalytic residues, including the native identities and all possible single amino acid variants (SAVs) at those positions. Linear fits and correlation coefficients are shown for each site. **(C)** Comparison of mutation-induced changes in frustration ($\Delta$ frustration, mutant minus native) predicted by FrustrAI-Seq and FrustratometeR across all catalytic residues and SAVs. Triangle entries mark wildtype residues. Points are colored by mutated amino acid identity. **(D)** Mutational frustration landscape for catalytic residues, showing raw frustration values and the delta of mutation-induced changes (FrustratometeR minus FrustrAI-Seq) summarized as heatmaps for each SAV. Boxed entries indicate native residues.

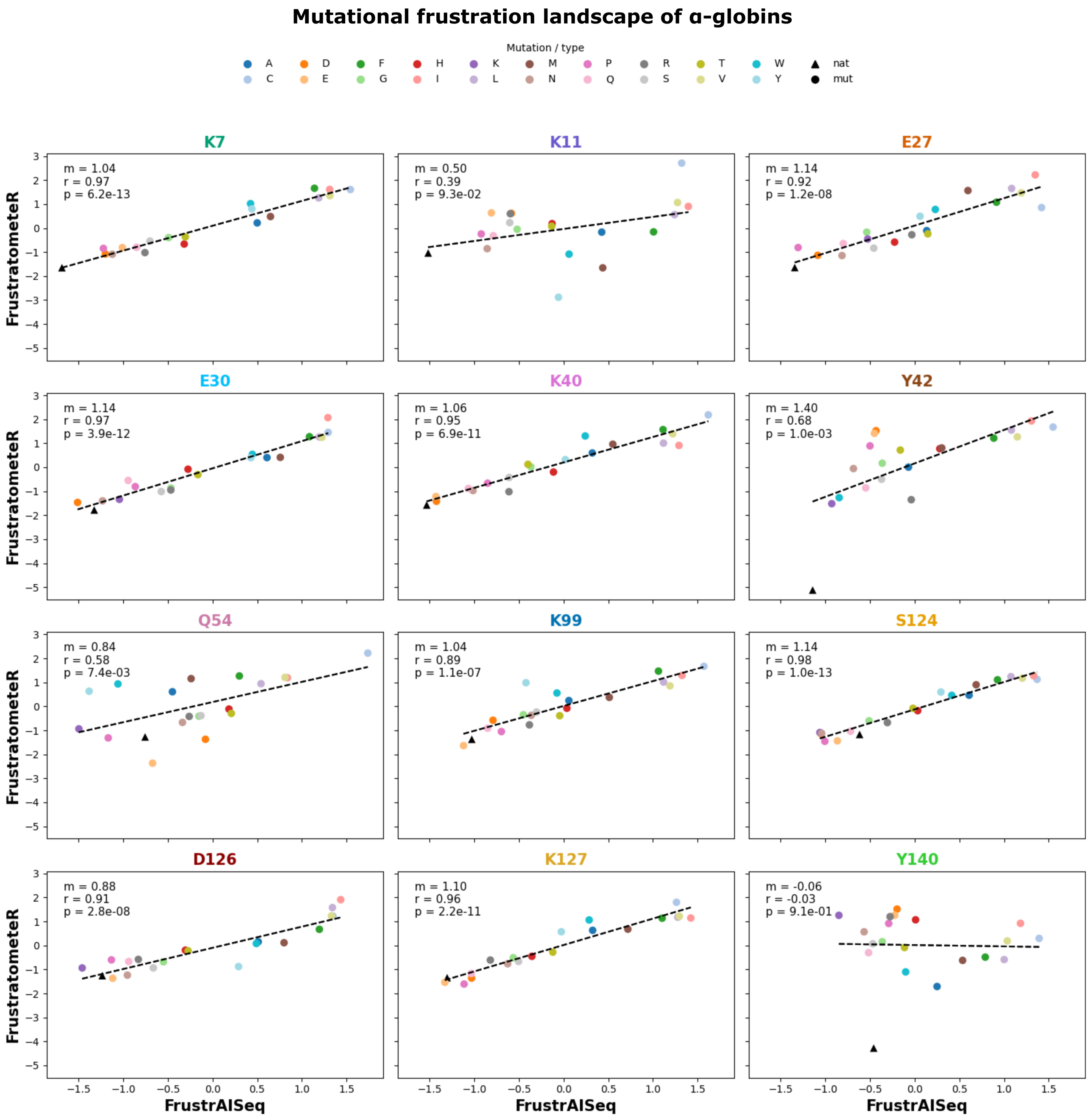


**Supplementary Figure 7. Mutational frustration landscape for positions in $\alpha$-globin structure.** Residues shown are natively conserved and involved in PPIs. Deltas of mutation-induced changes (mutant - wildtype) are shown for each residue and SAV. Triangle entries indicate native residues. Linear fits and correlation coefficients are shown for each site.

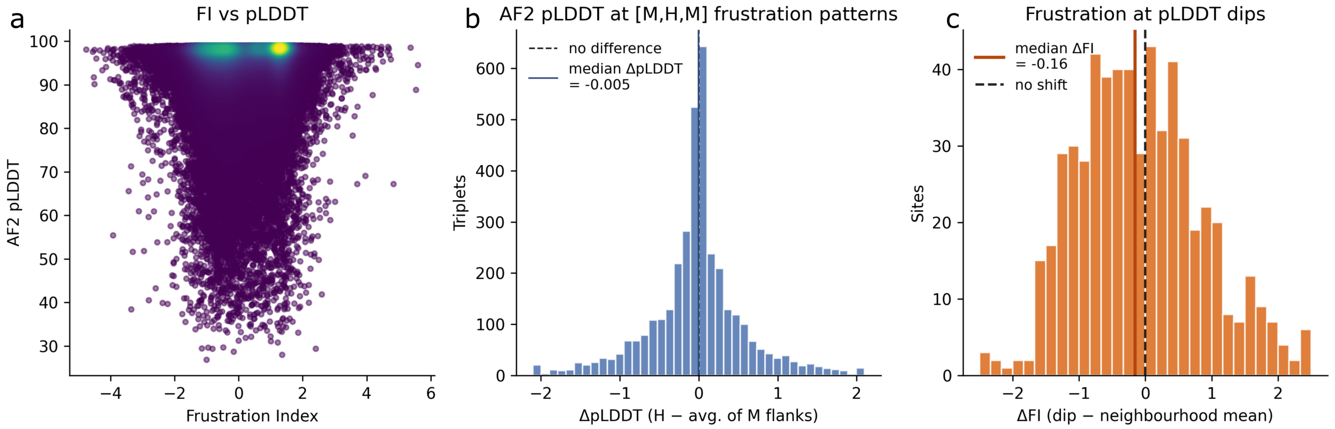


**Supplementary Figure 8. AlphaFold2 confidence (pLDDT) does not track local energetic frustration in high-quality AlphaFold2 structures (median pLDDT $\geq 90$).** Analysis on a random sample of 1000 Funstration test-set proteins for which per-residue AlphaFold2 pLDDT was retrieved from the AlphaFold Database and compared against FrustratometeR per-residue frustration indices. **(a)** Global association between FI and pLDDT across all residues (colored by point density). The two are only weakly related (Spearman $\rho = 0.12$), and the median pLDDT is essentially identical across frustration states (97.19, 97.06, and 97.88 for highly frustrated, neutral, and minimally frustrated residues, respectively). **(b)** Frustration-anchored test. For every $[\mathrm{M}, \mathrm{H}, \mathrm{M}]$ triplet (a highly frustrated residue flanked by two minimally frustrated residues), the pLDDT difference between the central frustrated residue and the mean of its two flanks ($\Delta$pLDDT). The distribution is centred on zero (median $-0.005$, $n = 3829$ triplets; Wilcoxon signed-rank $p = 1.1 \times 10^{-3}$): frustrated residues receive no lower confidence than their non-frustrated neighbors. **(c)** Structure-anchored test. At pronounced pLDDT dips (a residue whose pLDDT is $\geq 10$ below the mean of its $\pm 3$ flanking residues on both sides), the FI difference between the dip residue and the mean FI of its six flanking residues ($\Delta$FI). Dips coincide with only a marginal increase in frustration (median $-0.16$, $n = 585$; Wilcoxon signed-rank $p = 9.0 \times 10^{-3}$). Dashed lines mark zero difference, solid lines the observed medians. In both (b) and (c), the effects are negligible in absolute terms, and significance could be due to large sample sizes. Together, these analyses indicate that in the Funstration test set (high-confidence filtered: per protein median pLDDT $\geq 90$) there is no observable connection from low AlphaFold2 pLDDT towards any of the three frustration states.

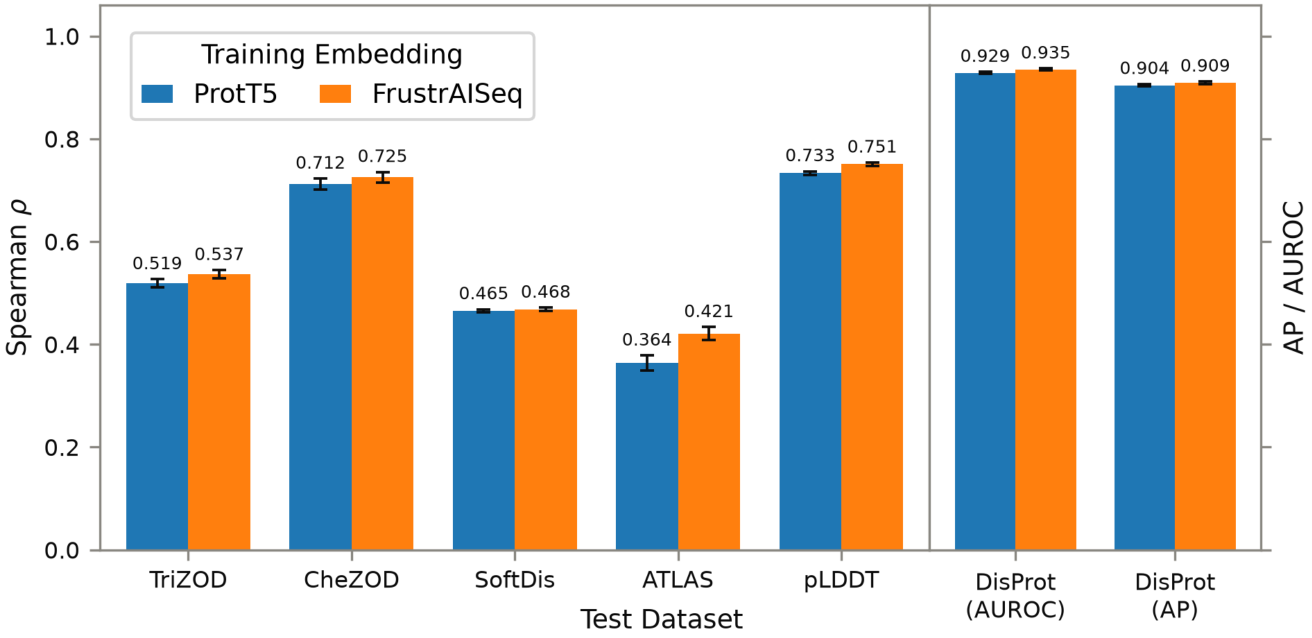


**Supplementary Figure 9. Prediction of intrinsic disorder when training on ProtT5 or FrustrAI-Seq embeddings.** To assess the effect of frustration finetuning on disorder captured by embeddings, a neural network (FNN with two hidden layers, 256-d and 64-d; LeakyReLU, dropout of 0.4, batch size 64, AdamW optimizer, MSE loss) was trained to predict continuous disorder as defined by TriZOD G-scores (1) (n=1027 proteins) either from a) ProtT5 (2) or b) FrustrAI-Seq embeddings as input. Evaluation was performed on six different datasets of protein intrinsic disorder and flexibility akin to the UdonPred pipeline (3):

**TriZOD** G-scores (n=348) and **CheZOD** (4) Z-scores (n=117), both derived from NMR chemical shifts; intrinsically and soft disordered regions from **SoftDis** (5), derived from missing coordinates in X-ray crystal structures and B-factors, respectively (n=5044); the **ATLAS** database (6) of Root Mean Square Fluctuation (RMSF) values derived from all-atom molecular dynamics simulations for representative protein structures from the PDB (7) (n=450); predicted local distance difference test (pLDDT) scores from AlphaFoldDB (8) (n=626), which strongly correlate with intrinsic disorder (9); and binary annotations from the DisProt (10) database, specifically the *Disorder-PDB* set of the CAID3 challenge (11) (n=348).

**Left panel** shows Spearman's rank correlation coefficient ($\rho$) between predicted scores and reference labels on continuous benchmarks. For CheZOD and pLDDT, whose label conventions increase with order rather than disorder, the reference labels were sign-inverted prior to correlation so that a larger $\rho$ always denotes closer agreement with the ground truth. **Right panel** shows performance on the binary DisProt disorder annotations, quantified as the area under the receiver-operating-characteristic curve (AUROC) and the average precision (AP). Predicted scores are pooled across all residues carrying a defined label and Gaussian-smoothed along the sequence ($\sigma = 1.5$ residues). Bars show the mean computed over the full pooled residue set, error bars denote the 95% confidence interval ($1.96 \times SEM$) across 100 bootstrap resamples of the pooled per-residue predictions. The TriZOD training set as well as the three non-fixed test sets (SoftDis, ATLAS, pLDDT - TriZOD, CheZOD, and DisProt have fixed test sets) were split such that no sequences between train and test had more than 30% sequence identity at 80% coverage (3). Compared to the evaluation in UdonPred, the PDBflex test set is omitted because both embeddings yielded correlations statistically indistinguishable from zero on it.

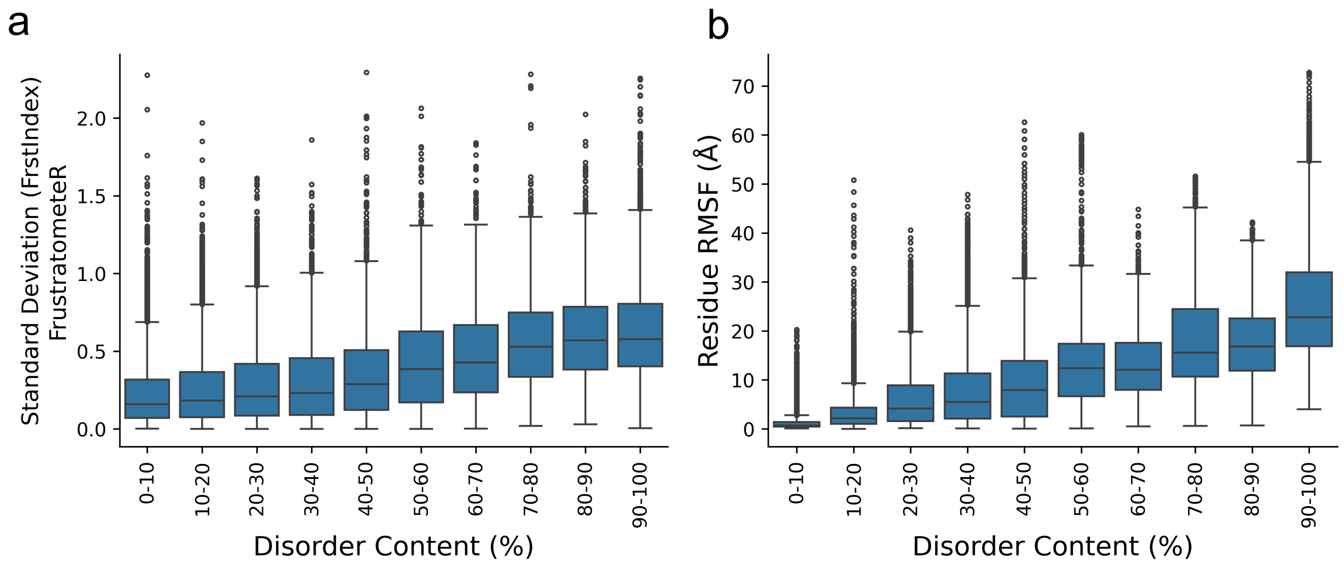


**Supplementary Figure 10. Conformational flexibility and frustration variability in disordered protein ensembles. a)** Distribution of the per-residue variability in frustration across conformers, stratified by protein disorder content. For each protein ensemble, the standard deviation of the FrustratometeR frustration values was calculated independently for each residue across all conformers in that ensemble. **b)** Distribution of per-residue root-mean-square fluctuations (RMSF) across conformers for the same protein ensembles, stratified by disorder content. Each point represents one residue. Before RMSF calculation, all conformers within each ensemble were structurally aligned to the first conformer using the `MDAnalysis.analysis.align` module. In both panels, each point represents one residue.

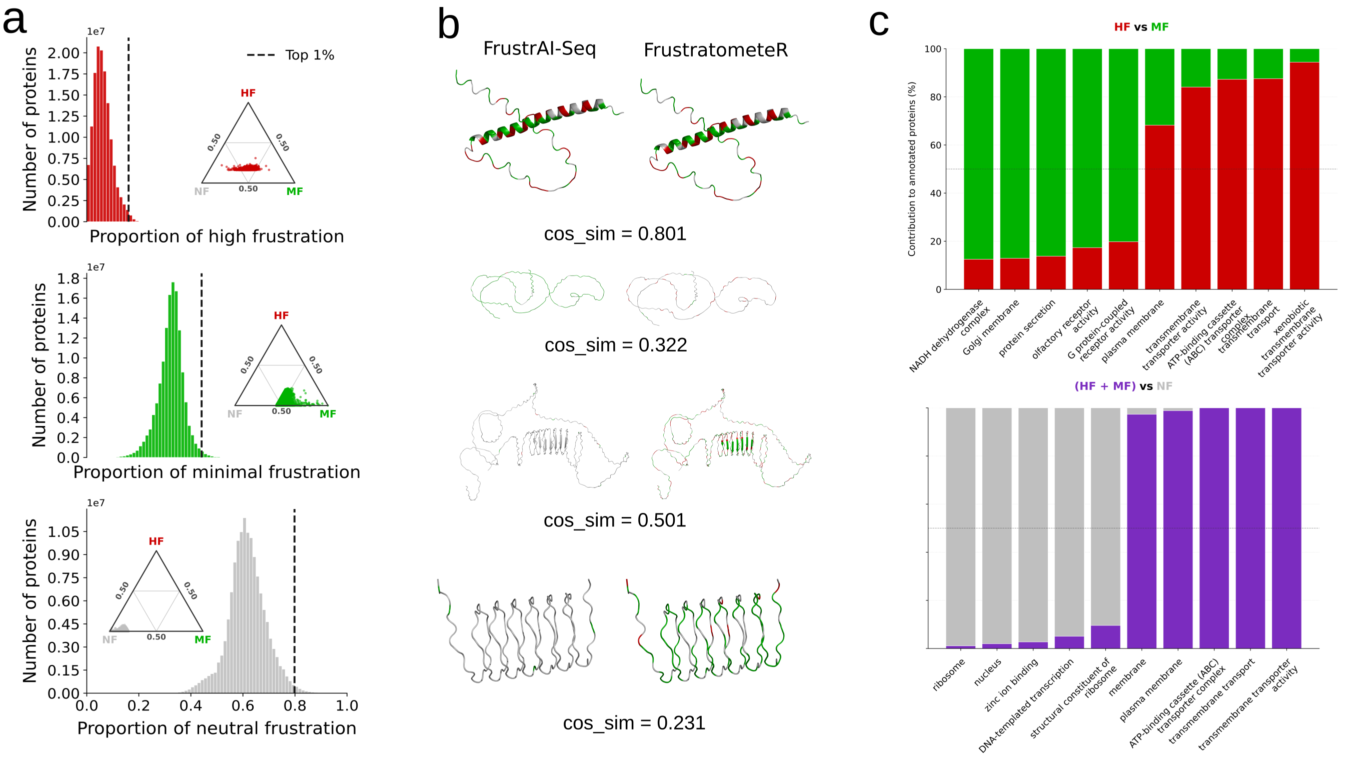


**Supplementary Figure 11. Proteome-scale comparison of frustration composition predicted by FrustrAI-Seq and calculated with FrustratometeR. a)** Distributions of the per-protein proportions of highly frustrated (top), minimally frustrated (middle), and neutral residues (bottom) across 255,000 proteins from 22,000 proteomes supported by both protein- and transcript-level evidence. Dashed vertical lines indicate the top 1% of proteins for each frustration class. Insets show ternary representations of the same top 1% proteins according to their relative proportions of highly frustrated (HF), minimally frustrated (MF), and neutral (NF) residues. **b)** Example proteins selected from the top 1% of each frustration class, comparing residue-level assignments from FrustrAI-Seq (left) and FrustratometeR (right). Residues are colored according to frustration class: minimally frustrated in green, neutral in gray, and highly frustrated in red. Cosine similarity values quantify the agreement between the two residue-level assignments. Examples with low cosine similarity (middle and bottom) are almost entirely homo-polymeric protein sequences. **c)** Gene Ontology term comparisons among proteins with extreme frustration compositions. The upper panel compares enriched terms associated with proteins dominated by highly frustrated versus minimally frustrated residues, whereas the lower panel compares proteins enriched in highly and minimally frustrated residues with those dominated by neutral residues. Bar heights indicate the relative contribution of each protein group to the annotated term.
